# Language Model Exploration and Survival Behavior Under Offline Replay of Organoid Neural Signals

**DOI:** 10.64898/2026.11.26.769443

**Authors:** Haoman Chen, Qingzu He, Fanxuan Chen, Xinyu Chen, Yang Liu, Pengcheng Yan, Jianwei Shuai

## Abstract

We tested whether replaying organoid signals recorded with a microelectrode array changes how a language model explores and survives in a game. In two trials on chip 22021, each group used its assigned readout during both training and evaluation, with zero additional reward weight. In the first trial, 16 training seeds per group were evaluated on 48 shared maps. After six decisions, the group receiving spontaneous recordings obtained before stimulation revealed 1.99 more cells per game than the all-zero group. The group receiving recordings from unstimulated trials revealed 2.16 more cells. The respective 95% intervals from resampling seeds on these fixed maps were [1.18, 2.73] and [1.58, 2.75]. Both median paired differences were zero. The two groups in each comparison revealed equal numbers of cells in 615/768 and 602/768 seed-map pairs, respectively. At equal elapsed time (game tick 6), the mean differences remained positive, but the 98.333% sensitivity intervals that varied both seeds and maps crossed zero. In the second trial, 32 seeds per group were evaluated on 64 reused pilot maps. We compared ordered replay with all-zero and block-shuffled inputs for exploration from low health and survival from severe thirst or hunger. All six adjusted intervals crossed zero. Earlier copies of 99 of 18,720 relabeled receipts were available, leaving the original protocol lock uncertain. On a separate chip, all three intervals for six-decision exploration crossed zero. On chip 22021, comparisons of post-stimulation and unstimulated recordings also yielded intervals crossing zero for exploration and score. These data show a fixed-map exploration difference after six decisions. Generalization to new maps and the separate contributions of training inputs, inference-time readouts, and the organoid source remain unresolved.

## 1 Introduction

Cultured neural tissue responds to electrical stimulation [1]. The challenge is to isolate the tissue’s own contribution to behavioral outcomes.

The model used Qwen3.5-0.8B [2], with approximately eight hundred million parameters, to select actions in Yaye. The four-dimensional electrode readings were included in the policy’s textual observations and fed into the value network.

We separately compared the number of newly revealed cells after six decisions and the number revealed through game tick 6 in the same trajectories. The first comparison fixed the number of choices, and the second fixed the elapsed game time.

This paper reports only changes in model behavior under specific inputs. A separate experiment is needed to determine whether the organoid learned during gameplay.

## 2 Related Work

Kagan et al. connected neurons cultured in vitro to a simulated game. Electrodes conveyed the game state to the neurons, and readout signals controlled the paddle [1]. Robbins et al. coupled mouse cortical organoids to CartPole and compared reinforcement-learning-selected stimulation with random stimulation and no stimulation [3]. In this study, a language model selected actions and received game feedback and prerecorded electrode observations. The study estimated game performance under these offline replay conditions and did not measure learning by the tissue during the game.

Organoid intelligence research treats three-dimensional tissue, electrodes, and computing systems as interrelated objects of measurement [4]. Cai et al. used a brain organoid connected to a high-density electrode array as a reservoir for speech recognition and nonlinear equation prediction [5].

Voyager used an automated curriculum, reusable skill code, and environmental feedback to advance exploration in Minecraft [6]. We measured exploration by counting newly revealed cells.

Fortunato et al. added learnable noise to network weights and examined changes in exploration across multiple game tasks [7].

## 3 Methods

Offline replay of the electrode recordings followed a separate path and a predetermined schedule. The training implementation used proximal policy optimization (PPO) [8] and updated low-rank policy adapters [9] and a separate value network.

The six-decision trial using recordings from the first organoid recorded on chip 22021 compared the intact group, the unstimulated group, and the all-zero group. It had no shuffled group. Each seed in each group was evaluated on the same 48 maps. The adverse-start trial using recordings from the second organoid recorded on the same chip compared the intact group, the all-zero group, and the shuffled group with two-second blocks shuffled within trials. For each group and scenario, each of the 32 training seeds was evaluated on the same 64 maps. In both training and evaluation, the two trials on chip 22021 used each group’s respective input scheme. The six-decision trial on the separate chip 22554 included a native R1-only group, an intact group with ordered training pairings, and a shuffled group with shuffled training-trial pairings. The two groups that received recording input both preserved the order of pre- and post-stimulation recordings within any one trial and used the same ordered recordings for evaluation.

For chip 22021, the all-zero group set the four-dimensional neural inputs to the policy and value networks to zero during both training and evaluation. In the six-decision evaluation of the first organoid recorded on chip 22021, with 16 seeds *×* 48 maps per group, the intact group used recordings from before the session’s first nonzero stimulation pulse onset marker. The unstimulated group used measured recordings from physical trials 15 and 18. Neither of the two trials had any nonzero stimulation output.

The adverse-start trial using recordings from the second organoid recorded on chip 22021 was checked against the available written and machine registration drafts. Each training seed corresponded to three models, one for the intact group, one for the all-zero group, and one for the shuffled group with two-second blocks reordered within trials. During evaluation, model parameters were frozen, each group retained its own input scheme, and runs began in the same scenarios and on the same maps.

Each of the 96 frozen models was evaluated under 3 starting conditions, on 64 maps per condition, for a total of 18,432 games. For the low-health start, the primary endpoint was the number of newly revealed cells at game tick 60 or at death if death occurred earlier. For the severe-thirst and severe-hunger starts, the primary endpoint was the survival rate at game tick 60 in each case. The six primary comparisons subtracted the allzero group and the shuffled group, respectively, from the intact group.

### Ethics and stem cell provenance

The use of hPSCs was approved by the Ethics Committee of the Institutes of Biomedical Sciences at Fudan University, Shanghai, China (no. 28). The human embryonic stem cell line H9 (WA09) was provided by Prof. Su-Chun Zhang, Waisman Center, Madison, USA.

## 4 Experimental Setup

MaxLab sampled voltage at 20 kHz. For each specimen, the controller calculated a two-hour operating window from the start time in the software binding record. The time when the tissue was physically connected to the array remains to be verified.

For the source of the human embryonic stem cell line H9 (WA09), see the ethics statement in Methods.

Each of the three warm-up check windows for the first 22021 specimen screened the same 322 channels from one batch. Recalculation from the archived raw voltage windows yielded 243, 241, and 230 passing channels, respectively (75.5%, 74.8%, and 71.4%). All were below the original real-time training threshold of 95% per window. Training that day used offline replay after acquisition. The reason for the on-site switch remains to be verified.

On each of the two days, the run logs for 22021 listed the electrode coating as platinum black. The material certificate for the chip lot still needs to be verified. Table 3 on page 2 of the manufacturer’s guide [10] lists the recommended starting parameters for standard MaxOne/6WP platinum-black electrodes as 300 to 600 mV, with a phase width of 200 µs and a pulse interval of at least 300 ms. On each of the two days, biphasic commands were sent with the positive phase first, at 199.79 mV and 200 µs per phase. In the regular condition, successive pulse onsets on the same electrode were 200 ms apart, based on the device commands and event markers. The basis for converting the guide’s amplitudes to per-phase DAC commands still needs to be verified. The actual waveform and peak amplitude under electrode load were not measured on site, and actual tissue exposure still cannot be determined.

On 11/18, the validation receipt for the fixed readout panel of 22554 recorded a validation failure. For the exploratory acquisition that followed, the startup check confirmed the panel validation result before the first trial, and the acquisition was logged as not admitted in the acquisition protocol’s real-time admission field. Each block contained one trial for each of H, W, and F, with 3 trials per site in total. Only after acquisition ended was exploratory leave-one-block-out classification performed separately on the pre- and post-stimulation recordings, pooling all 9 trials. Both pre- and post-stimulation classification yielded 5/9 correct predictions, with 1/3, 3/3, and 1/3 correct for H, W, and F, respectively. The exact permutation results were p=0.25 before stimulation and p=0.30556 (approximately 0.306) after stimulation. The reference criteria for the qualification procedure were at least 7/9 correct after stimulation, at least 2/3 correct at each site, post-stimulation p*≤*0.05, and pre-stimulation background p>0.05. The post-stimulation result did not meet these criteria, and the qualification procedure was not invoked for this exploratory acquisition.

According to the accompanying per-model training budget manifest (per_model.csv), the initial and expansion cohorts in each of the two six-decision trials each contained 8 models per group. Each model completed 4 collection batches of 12 decisions each, for a total of 48 training decisions, 8 policy optimization steps, and 64 value optimization steps.

For the archived six-decision intervals, the mean paired difference was first calculated for each training seed over a fixed set of 48 maps. Training seeds were then resampled with replacement 12,000 times to obtain two-sided 95% percentile intervals. For the two primary six-decision comparisons on the first specimen of 22021, the archived intervals used the original fixed set of 48 maps and resampled from the 16 seeds pooled across the initial and expansion batches. The proportions of the two batches could vary across resamples. The post hoc sensitivity analysis held fixed the original 48 maps and the allocation of 8 seeds per batch. It sampled 8 seeds with replacement within each of the initial and expansion batches, then pooled the seed-level mean differences from the two batches with equal weights. Exact convolution of the integer-valued distribution of total paired differences in counts of newly revealed cells yielded two-sided 95% percentile intervals.

## 5 Results

For the first specimen on chip 22021, we compared the count of newly revealed cells after six decisions per episode across 16 seeds and 48 maps. The all-zero group, intact group, and unstimulated group each had 768 episodes. Relative to the all-zero group, the intact group and the unstimulated group revealed 1.99 and 2.16 more cells per episode on average, respectively. The post hoc analysis held the original 48 maps fixed, resampled 8 seeds within each of the two cohorts, and pooled the cohorts with equal weights. The two-sided 95% intervals for the intact group minus the all-zero group and the unstimulated group minus the all-zero group were [1.204, 2.742] and [1.590, 2.707] cells per episode, respectively. Both intervals were entirely above zero. Inference was limited to this record, the original 48 maps, and a composition in which each of the two cohorts contributed half.

The intact group and the unstimulated group were each paired with the all-zero group by training seed and map, yielding 768 episode pairs per comparison. In that order, the counts of increases, ties, and decreases in newly revealed cells were 101/615/52 and 109/602/57 for the two comparisons. The median paired difference was 0 cells in both of the two comparisons.

Readouts following the two stimulation conditions, regular and random, were each paired with readouts from the unstimulated group. On 11/18, the differences in newly revealed cells were *−*0.8125 [*−*3.182, 0.906] and +0.1953 [*−*1.495, 1.682] cells, respectively, and the R1 differences were *−*0.000777 [*−*0.003446, 0.001326] and +0.000576 [*−*0.001536, 0.003840] points, respectively. On 11/19, the differences in newly revealed cells were *−*0.4870 [*−*2.760, 1.424] and *−*0.2865 [*−*2.706, 2.401] cells, respectively, and the R1 differences were *−*0.000785 [*−*0.003990, 0.001539] and *−*0.000738 [*−*0.004779, 0.002393] points, respectively. All eight of these 99.167% intervals crossed zero.

For 22554, the six-decision evaluation used recordings from 11/18. The critic_input_size field read directly from the evaluation logs indicated that the native R1-only group did not append the four-dimensional MEA readings and had a 108-dimensional value network input. The intact group (ordered training pairings) and the shuffled group (shuffled pairings of training trials) each had a 112-dimensional input. All intervals for the exploration differences crossed zero, leaving the effects of training pairings or input conditions on exploration inconclusive.

The severe-thirst start began with 1.6 hydration points, and the severe-hunger start began with 4 satiety points. In the two scenarios, the point estimates of the survival-rate differences for the intact group minus each of the two control groups were all negative (Table 2). All four full-sample 99.167% intervals crossed zero. For the intact group minus the all-zero group and minus the shuffled group, in that order, the upper bounds were +2.73 and +2.83 percentage points under severe thirst and +2.34 and +3.29 percentage points under severe hunger. Smaller improvements and decreases in survival rate remained compatible with these intervals. A review against the available registration draft on all 64 reused pilot maps, including the analysis of exploration under the low-health start, found that none of the six primary comparisons met the criteria (0/6) and none of the three scenarios met the criteria (0/3).

## 6 Analysis

In a post hoc exploratory analysis of the same six-decision trajectories from Section S5.1, we replayed the recorded actions and truncated the replay at game tick 6. In paired comparisons with the all-zero group, the intact group revealed 1.21 more cells and the unstimulated group revealed 1.53 more cells. The 95% intervals from crossed resampling of seeds and maps [11] were [0.08, 2.59] and [0.14, 3.18], respectively. A post hoc sensitivity analysis used the nominal Bonferroni level of 98.333% for the three nonzero *β* comparisons in the original *β* sweep. The two archived crossed-resampling intervals were [*−*0.16, 2.97] and [*−*0.12, 3.65] cells per episode, respectively. Both crossed zero. When the estimand allowed the maps to vary, the exploration difference at game tick 6 remained inconclusive.

At game tick 6, in the same set of 768 trajectory pairs, the win/tie/loss counts were 79/654/35 for the intact group minus the all-zero group and 88/644/36 for the unstimulated group minus the all-zero group. When only the 38 largest positive differences were removed, the mean differences across the remaining 730 pairs were *−*0.138 and +0.095 cells.

## 7 Discussion

On chip 22021, each group used its own input-reading procedure for both training and evaluation in the six-decision experiment with the first specimen and in the stress trial with the second specimen. Paired differences for the same seed and map cannot distinguish action preferences acquired during training from the frozen model’s response to the current step’s readings at inference. Setting the model’s neural input to zero in the all-zero group simultaneously removed differences in magnitude, variability, correlation structure, and temporal variation in the four readings. Generic time-varying inputs could still explain the observed group differences. The existing comparisons cannot isolate a contribution specific to the organoid source. The first-step diagnostic in Section S5.5 was limited to the outputs, under shared ordered replay, of the policies obtained from the two training pairings on chip 22554.

## 8 Limitations

On each of the two days on chip 22021, one organoid was recorded, and each organoid had only one acquisition session. Variation across independent organoids, chips, and sessions cannot yet be estimated separately.

On 11/19, sites H, W, and F on chip 22554 each had one screening stimulation recording for comparison of the preand post-stimulation windows. Same-site zero-output controls were unavailable. The separability of neural responses by stimulation site and the actual waveform under electrode load remain undetermined.

The written and machine-readable registration drafts for the adverse-start trials specified exploration for the low-health start and survival for the starts with near-fatal thirst and near-fatal hunger. The machine-readable registration draft was labeled “draft.” The design used for execution retained the legacy labels specifying survival assessment in all three scenarios and 6/8 seeds. The registration hash in the training design was a place-holder. The archived analysis code took the ceiling of 75% of the current seed count, giving a favorable-seed threshold of 24/32 for the full sample. The execution order of the three arms was rotated by seed and round. The frozen audit game source specified in the registration was missing. Execution used a frozen copy of the existing paired project. Cutoff observations were read using the Python adapter’s mapping of the old protocol’s landmark to observe together with the per-tick trajectories. The scenario verification file regenerated during execution had a hash that differed from the old registered hash. The older file recovered afterward matched the old registered value. After evaluation, the fingerprint was updated, as were the design identifiers in 18,720 receipts. These comprised 18,432 per-game receipts and 288 evaluation freeze receipts. Only 99 receipts could be compared individually against independent earlier copies (98 per-game receipts and 1 evaluation freeze receipt). After replacing only the design identifier, the before and after versions of those 99 receipts were byte-for-byte identical. All 98 comparable per-game receipts came from the 8 seeds that overlapped with the pilot. The remaining 18,621 receipts lack corresponding earlier copies, including 18,334 per-game receipts and 287 evaluation freeze receipts. The pre-unblinding state of these receipts cannot be verified individually. The eight-seed pilot documentation already specified that at least 6/8 seeds had to be favorable, and the 32-seed draft specified at least 75% favorable seeds. The archived copy of the machinereadable draft is byte-for-byte identical to the current machine-readable draft. The internal ordering indicated by archive member modification times is consistent with the draft predating training. The training design used for the run recorded a placeholder for the draft fingerprint, so that field cannot bind the run to this machine-readable draft. Whether the endpoints, thresholds, and run design had been uniquely fixed before unblinding, and whether all receipts had been sealed by then, still require independent verification. In Appendix B, the complete replay of saved actions and the spot checks that reloaded the final models checked the execution results. Neither can recover the pre-unblinding state of these receipts.

All 64 evaluation maps and 8 of the 32 training seeds had been used in the pilot. The post hoc sensitivity analysis retained only the remaining 24 seeds but still used all 64 pilot maps. Independent validation on new maps remained unavailable.

## 9 Conclusion

In the offline replay of recordings from the first organoid recorded on chip 22021, under the same budget of six decisions, the intact group and the unstimulated group revealed an average of 1.99 and 2.16 more cells per run than the all-zero group, respectively. When runs were paired by training seed and map, the median difference was zero in each of the two comparisons.

The second organoid recorded on chip 22021 was reassessed on all 64 reused pilot maps according to the available registration draft. None of the six primary comparisons met the criteria (0/6). Subjective fear was not measured.

## Technical appendix

### Extended summary

We used a microelectrode array (MEA) to record organoid activity and used offline replay to feed the recordings to a language model that controlled a character in a survival game. For chip 22021, the six-decision trial on the first specimen and the adverse-start trial on the second specimen both used each group’s readout method during training and evaluation, with the additional reward weight set to zero. In the first specimen’s trial, the intact group (baseline, archived figure label “self-generated”) used recordings of spontaneous activity from before the first stimulation. The unstimulated group (sham, archived figure label “zero-output”) used recordings measured in the corresponding round, when no nonzero stimulation was delivered. Prior stimulation may have affected those recordings. The all-zero group (masked, archived figure label “masked”) set the four-dimensional readout to zero. This trial had no shuffled group. Each of the 16 training seeds per group was evaluated on the same 48 maps. After six decisions, the intact group and the unstimulated group revealed an average of 1.99 and 2.16 more cells per game than the all-zero group, respectively. The 95% intervals from resampling training seeds on the fixed map set were [1.18, 2.73] and [1.58, 2.75], respectively. When runs were paired by training seed and map, the median was zero for each of the two differences. The counts of newly revealed cells were equal in 615/768 and 602/768 pairs, respectively, and were identical across all three groups in 577/768 seed–map combinations. When the same trajectories were truncated post hoc at game tick 6, the two mean differences remained positive. In a sensitivity analysis using the nominal 98.333% level from the original reward-weight sweep, the two intervals for the fixed map set were entirely above zero, whereas the intervals from crossed resampling of seeds and maps all crossed zero. The difference in exploration at equal elapsed time remained uncertain on different maps. Subtracting the all-zero group’s shares from the unstimulated group’s shares for the actions actually selected gave differences of *−*2.69, +2.28, and +1.22 percentage points for single-cell movement, landmark navigation, and gathering, respectively, with 4,608 decisions per group. In the first-day five-group subexperiment, the intact group and the unstimulated group were each compared with the all-zero group using 384 pairs per comparison. The timing component of the score associated with action duration accounted for 84.3% and 95.9% of the two total score differences, respectively. The trial on the second specimen included an intact group with ordered replay, an all-zero group, and a shuffled group in which two-second blocks were shuffled within each trial. The evaluation used 32 seeds *×* 64 maps for each group under each start condition. Following the available registration draft, we rechecked the endpoints on all 64 reused pilot maps. We assessed exploration in the low-health start through game tick 60 or until death if it occurred earlier. We also assessed survival at game tick 60 for the severe-thirst and severe-hunger starts. For each of the three endpoints, we compared the intact group against the all-zero group and against the shuffled group. The adjusted intervals for all six primary comparisons crossed zero, and none met the criterion (0/6). Of the 18,720 receipts relabeled after evaluation, 99 had independent earlier copies that could be used to check the scope of relabeling. These covered 3 of the 9 scenario–group combinations. All 98 per-game receipts in this subset came from pilot seeds, and the subset also included 1 evaluation-freeze receipt. Independent verification is still needed to establish whether a single protocol was locked before unblinding, which protocol it was, and whether the full set was sealed. For chip 22554, we separately compared the native R1-only group, the intact group with ordered training pairings, and the shuffled group with shuffled training-trial pairings. The two groups receiving MEA input used the same ordered recordings during evaluation. All three intervals for exploration after six decisions crossed zero. For chip 22021, across two days, we subtracted the unstimulated group’s outcomes from those of the post-regular-stimulation group and, separately, from those of the post-random-stimulation group. The intervals for exploration and score all crossed zero. The existing design for chip 22021 did not allow the specific contributions of the training readout method, the instantaneous readouts during inference, and the organoid source to be identified separately.

The technical appendix uses Sections S1–S9. Figure, table, and algorithm numbers run through the main article and its appendices.

## S1 Introduction

Cultured neural tissue responds to electrical stimulation [1]. The challenge is to isolate the tissue’s own contribution to behavioral outcomes. DishBrain connected neurons to a simulated table tennis game. Neural firing directly controlled the paddle, and performance changed in turn [1]. The pathway was short, but more than one explanation was possible. The tissue may have exhibited plasticity. Any time-varying external input might also have kept the policy from repeatedly selecting the same few actions.

The model used Qwen3.5-0.8B [2], with approximately eight hundred million parameters, to select actions in Yaye. The four-dimensional electrode readings were included in the policy’s textual observations and fed into the value network. For chip 22021, both the six-decision trial with the first specimen and the adverse-start trial with the second specimen used the game score R1 as the training reward. The first-day six-decision trial for chip 22554 used the same training reward, and the additional reward weight *β* was zero in all these trials. In the second-day trial for chip 22554, which held out data by recording-site file, the two groups that received electrode input also used an auxiliary reward assigned according to replay-window phase. Appendix B describes the training and evaluation procedures. In each trial, the readings could indirectly change the game score through action selection. For chip 22021, training and evaluation both followed each group’s own readout method. The between-group differences therefore encompassed action preferences learned during training and responses to the readings at inference time.

The game made the differences measurable. In the short six-decision trial, six action selections were evaluated per episode. The character could retrace its previous route or navigate toward a landmark, revealing new cells along the way. Actions such as gathering and waiting also counted toward the decision budget. Yaye recorded the number of newly seen cells in each episode, counting each cell only one time. With the starting point, map, and decision budget fixed, how many more cells were seen was a direct measure that required no subjective judgment.

We separately compared the number of newly revealed cells after six decisions and the number revealed through game tick 6 in the same trajectories. The first comparison fixed the number of choices, and the second fixed the elapsed game time. We separately broke down the score by how many game ticks each action took. Action frequencies described the choices the model actually made. We tested the scope of the results separately using recordings from two days, one additional chip, and three types of adverse starts. We used the pilot matrix to estimate the probability of meeting the predefined criterion at the specified magnitude of improvement.

We state the scope of the study upfront. This paper reports only changes in model behavior under specific inputs. A separate experiment is needed to determine whether the organoid learned during gameplay.

## S2 Related Work

### S2.1 Closed-Loop Experiments with Cultured Neural Tissue

Organoid intelligence research treats three-dimensional tissue, electrodes, and computing systems as interrelated objects of measurement [4]. Cai et al. used a brain organoid connected to a high-density electrode array as a reservoir for speech recognition and nonlinear equation prediction [5]. Liu et al. encoded tactile inputs as stimulation and classified Braille letters from organoid responses in an open-loop task [12]. In the main comparison, the two acquisitions for 22021 came from two organoids on the same chip, and the recordings for 22554 came from one other chip. Training seeds reproduce random variation in model training, and test maps reproduce game conditions. Neither can increase the number of independent biological samples.

### S2.2 Measuring Exploration in Games

Voyager used an automated curriculum, reusable skill code, and environmental feedback to advance exploration in Minecraft [6]. In Yaye, we evaluated the actions each group selected on its own using the same starting points, retained maps, and decision budgets. We measured exploration by counting newly revealed cells. Pairing exposed each group to the same map conditions while allowing between-group differences to vary by map.

Actions such as navigation can span multiple game ticks, so the same number of decisions can correspond to different amounts of game time.

### S2.3 Input Perturbations and Exploration

Fortunato et al. added learnable noise to network weights and examined changes in exploration across multiple game tasks [7]. In this study, temporal variation came from the four-dimensional observations read by the model. If time-varying input in general is sufficient to explain the behavioral differences, synthetic input that matches the statistical properties of the measured signals could also produce similar results. We have not yet directly compared these two types of input.

To compare exploration performance under two evaluation budgets, the study reported paired results for a fixed number of decisions alongside paired results obtained by retrospectively truncating the same set of trajectories to a fixed number of game ticks. It used counts of ties by episode and by map to describe how the positive mean difference was distributed across the evaluated episodes. See Sections S5.1 and S6.

## S3 Methods

### S3.1 Game–model feedback loop and replay of electrode recordings

Yaye is a grid-based survival game. The character has health, hydration, and satiety levels and can move, navigate to a destination, gather, eat, drink, rest, or wait. The world advances in time steps (ticks). A valid move to an adjacent cell takes one game tick. Navigation advances according to the number of steps actually taken. In this experiment, each navigation action covered at most four cells and took correspondingly less time if it stopped early.

Figure 1 illustrates the offline neural input. The game state was fed back to the model, and actions selected by the model in turn changed the game state. Offline replay of the electrode recordings followed a separate path and a predetermined schedule. The tissue did not receive game actions, and the current action cannot alter spikes that have already been recorded. The game state, model sampling, and the observations read by each group could all change over time. The first specimen on chip 22021 did not meet the channel stability threshold for real-time training (Section S4.1).

**Figure 1.**
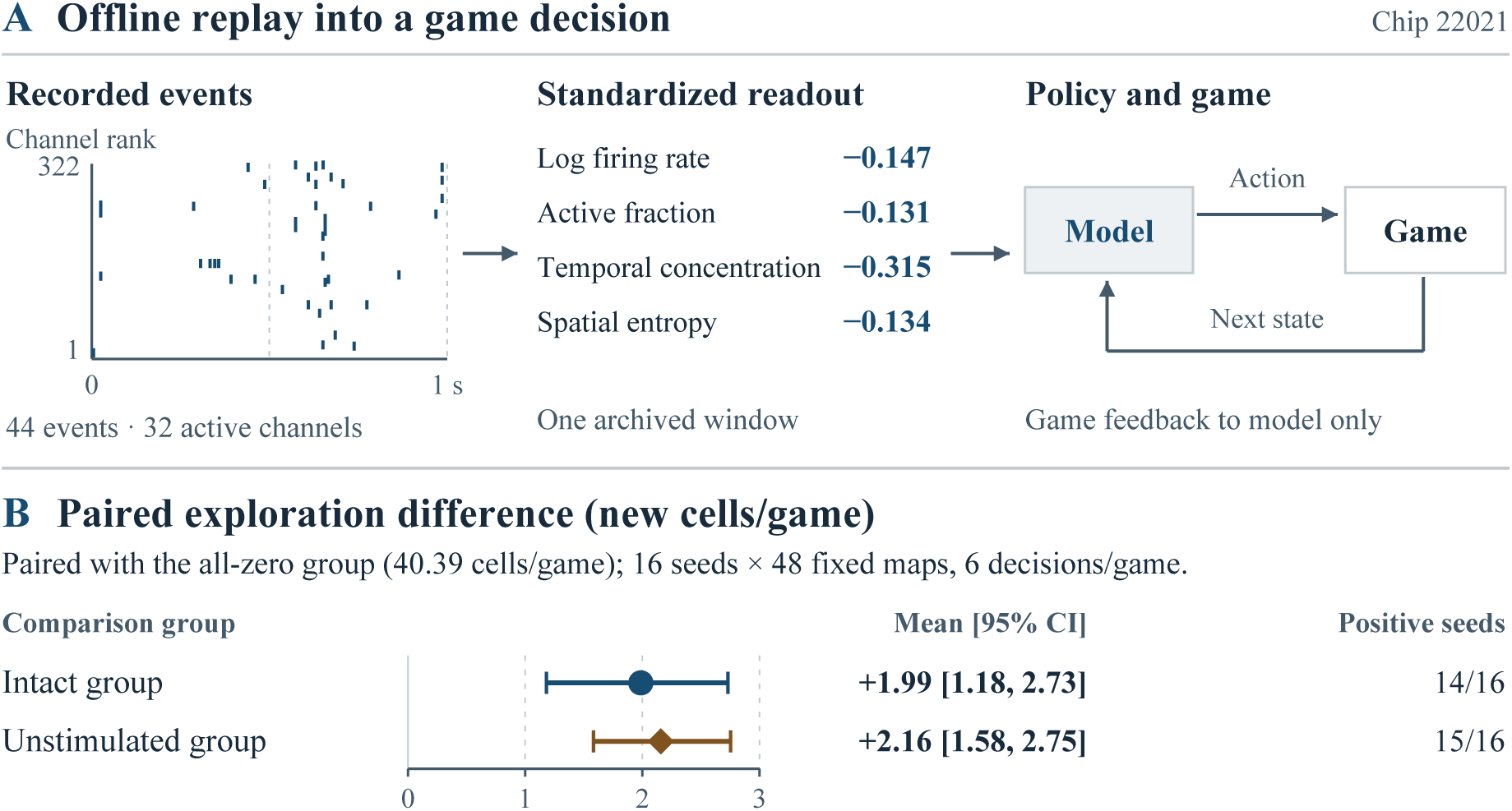
Offline neural input and exploration on chip 22021. (A) The recorded window covers [58, 59) seconds after warm-up began and precedes the session’s first nonzero pulse onset marker. The horizontal axis shows time in seconds relative to the start of the window, and the vertical axis shows the ranks of the selected channels in hardware-channel order. Each tick represents one recorded spike. During offline replay, the model received four standardized readouts, sent actions to the game, and received the resulting game states. (B) The intact group used pre-stimulation spontaneous activity, the unstimulated group used activity recorded while stimulation output was zero, and the all-zero group used constant-zero neural input. Each group used its assigned input in both training and evaluation. After six decisions per game, the intact and unstimulated groups were each compared with the all-zero group, which averaged 40.39 newly revealed cells per game. Points show the mean paired difference in newly revealed cells per game, calculated by subtracting the all-zero group’s result from the comparison group’s result. Lines show 95% confidence intervals obtained by resampling the 16 training seeds per group while holding the same 48 maps fixed. The Positive seeds column counts seeds for which the comparison group revealed more cells on average than the all-zero group across the 48 maps. The contrasts are exploratory and condition on this map set.

### S3.2 Construction of neural observations

At each step, the model connected to the MEA read the game state and the neural observation. The game state provided the surrounding terrain, reachable landmarks, inventory, and health, hydration, and satiety levels. In each one-second window, which included its left endpoint and excluded its right endpoint, N was the total spike count across the selected channels, C was the total number of selected channels, including silent channels, and A was the number of channels with spikes. The four dimensions were, in order, ln(1+N/C), A/C, 1*−*H_time/ln(10), and H_channel/ln(C). H_time was the Shannon entropy of the spike distribution across ten equal-width time bins, and H_channel was the Shannon entropy computed from each channel’s share of N. Both calculations used natural logarithms and skipped empty terms. All spikes were weighted equally. Before baseline standardization, the archived extraction program clipped temporal concentration and normalized channel entropy, each to the interval [0, 1]. In the training, end-of-batch allocation, and retained source windows checked in the two-day, five-group experiment, each of these two measures had the same value before and after clipping. For empty windows, all four raw dimensions were zero. The center for each dimension was the median across calibration windows, and the scale was the larger of 1.4826 times the median absolute deviation and a lower bound. The lower bound was 0.1 for the first dimension and 0.05 for each remaining dimension. For each dimension, the center was subtracted and the result was divided by the scale, clipped to [*−*3, 3], then divided by 3, with three decimal places retained. This vector was entered into the policy prompt and the value network. The all-zero group directly masked all four dimensions after calibration. The calibrated readout for an empty window could differ from zero.

The four-dimensional input was provided according to each group’s replay readout procedure. In one evaluation episode for the first specimen with training seed 102600001, all three groups used the same map and had the same encoding digest for the base game observation at the first step. The all-zero group read (0, 0, 0, 0), the intact group read (*−*0.147, *−*0.131, *−*0.315, *−*0.134), and the unstimulated group read (0.495, 0.502, 0.077, 0.230). Spike events were re-extracted from the retained warm-up window for the intact group and the trial window with zero stimulation output for the unstimulated group, both from the same session. The values recomputed using the formulas in this section matched the actual first-step readouts for these two groups component by component. The all-zero group referenced that same trial window with zero stimulation output, and its actual logged neural readout was zero. All three groups chose a single-cell move to the same destination cell. The log for that step did not retain the selection probabilities for individual action verbs or candidate targets. Each group continued to use its own readout procedure in training and evaluation. This episode could not distinguish the influence of preferences learned during training from that of the neural readout at that step. Section S7 compares the full set of replay conditions and describes the limits of attribution. The first-step probability diagnostic in Section S5.5 covers only the two training pairings under shared ordered replay on chip 22554.

### S3.3 Policy learning

The character was controlled by Qwen3.5-0.8B, and only a small number of additional parameters were updated during training. The project’s base-model manifest matches all 11 archived file hashes to the pinned upstream revision [2]. The hash comparison identifies the archived files. It cannot establish which files each historical training process loaded. The archived adapter configurations for all 48 policies in the first specimen’s six-decision analysis and all 96 adverse-start policies on chip 22021 specified low-rank adaptation [9], with rank 8, alpha 16, and dropout 0. The policy network selected actions, and the value network estimated future scores. The two networks were updated separately, and training used proximal policy optimization [8]. Both components received neural observations. When the input was all zero, the inputs to both components were set to zero together, so information cannot leak through the other branch.

### S3.4 Neural inputs and controls for each experiment

#### Replay groups and pairing budgets for each experiment

| Design and specimens | Manuscript group names (archived names) and input-reading procedures | Comparisons and evaluation budget |
|---|---|---|
| six decisions, first specimen on chip 22021 (11/18) | The intact group (baseline, archived figure label “self-generated”) used actual recordings collected before the first stimulation. The all-zero group (masked, archived figure label “masked”) set the four-dimensional readings for both the policy and value networks to zero. The unstimulated group (sham, archived figure label “zero-output”) used actual recordings made when there was no nonzero stimulation output in that round. Prior stimulation may have affected these recordings. | Training and evaluation both used each group’s input-reading procedure. The intact group and the unstimulated group were each compared with the all-zero group. The trial included three groups in total. Each of the 16 training seeds per group was evaluated on the same set of 48 maps, with six decisions per episode. Results at game tick 6 were obtained by truncating the same trajectories post hoc. |
| six decisions, chip 22554 (H/W/F record) | Native R1-only group (pure), intact group (ordered, ordered training pairings), and shuffled group (shuffled, shuffled training-trial pairings). | The trial used the 11/18 record. Each arm used 16 seeds × 48 maps, for a total of 768 episodes and 4,608 decision rows per arm. The fields in the evaluation logs that recorded the inputs actually read showed that the native R1-only group had no appended four-dimensional MEA input. Its value network had a 108-dimensional input, and each of the two groups with MEA input had a 112-dimensional value-network input. The two groups with MEA input differed in their training-trial pairings and used the same ordered four-dimensional input for evaluation. |
| five groups, chip 22021, one specimen on each of two days | The design retained the all-zero group (masked), intact group (baseline), and unstimulated group (sham), and added a group using recordings collected after regular stimulation (regular) and a group using recordings collected after random stimulation (irregular). | The groups using recordings collected after regular and random stimulation were each compared with the unstimulated group. The intact group and the unstimulated group were each compared with the all-zero group. The same 8 seeds × 48 maps were used on the two days, with 6 decisions per episode. |
| Adverse starts, second specimen on chip 22021 (11/19) | Intact group (ordered, replay in the original order), all-zero group (masked, readings masked), and shuffled group (shuffled, two-second blocks reordered within trials). | The intact group was compared separately with the all-zero group and the shuffled group. The shuffled group preserved the order within each block. The budget was 32 × 64 per arm per scenario. Readings were taken at game tick 60 or at death. |
| Holdout by site file, chip 22554 (11/19) | Intact group (ordered, ordered training pairings), shuffled group (shuffled, shuffled training donor pairings), and native R1-only group (pure, reuse of an existing checkpoint). | The intact group was compared separately with the native R1-only group and the shuffled group. Each group had 72 episodes, and readings at game ticks 48 and 192 came from the same trajectories. Appendix B describes holdout preprocessing and the training reward. |

The six-decision trial using recordings from the first organoid recorded on chip 22021 compared the intact group, the unstimulated group, and the all-zero group. It had no shuffled group. The initial and extended batches each had 8 training seeds, with no overlap between batches. Each seed in each group was evaluated on the same 48 maps. The archive can be used to verify the group-specific input schemes, evaluation budgets, and additional reward weights for the two batches. In the staging archive for the extended batch, baseline/train.py was the entry point for the intact group, and main/train.py was the entry point for the unstimulated group and the all-zero group. The two retained entry-point files differed only in the group names accepted on the command line, and each was byte-for-byte identical to its counterpart in the initial batch. The intact group’s entry point in the software package accompanying the manuscript also contains an auxiliary reward mixing branch and writes the reward_mix field. The field was absent from all 8 model registration records for the intact group in the extended batch. For every batch update, the receipts for these models recorded an actual auxiliary reward contribution of zero. The available receipts can be traced to each model’s input registration and update records. Runtime fingerprints are still missing for the entry point loaded by the actual process, the imported dependencies, and the full runtime environment. The adverse-start trial using recordings from the second organoid recorded on the same chip compared the intact group, the all-zero group, and the shuffled group with two-second blocks shuffled within trials. For each group and scenario, each of the 32 training seeds was evaluated on the same 64 maps. In both training and evaluation, the two trials on chip 22021 used each group’s respective input scheme. The shuffled group preserved order within each two-second block. A step-by-step check of the archive found that all training seeds in the shuffled group used the same sequence of training windows. The retained evaluations used map-specific block permutations. For a given map and game tick, the designated window was reused across training seeds and starting scenarios. Different maps could use different windows at the same game tick. Whether the models identified or made use of the maps through these assignments remains unknown. Reordering the two-second blocks changed both continuity across blocks and the correspondence between decisions and recording segments. The difference between the intact group and the shuffled group describes the performance difference under the replay arrangement used. The individual effects of the two changes still cannot be separated. The six-decision trial on the separate chip 22554 included a native R1-only group, an intact group with ordered training pairings, and a shuffled group with shuffled training-trial pairings. The two groups that received recording input both preserved the order of preand post-stimulation recordings within any one trial and used the same ordered recordings for evaluation. The group table in this section lists the recordings and budgets for each group.

For chip 22021, the all-zero group set the four-dimensional neural inputs to the policy and value networks to zero during both training and evaluation. This simultaneously removed the amplitude, variability, correlation structure, and temporal variation of the readouts. In the six-decision evaluation of the first organoid recorded on chip 22021, with 16 seeds *×* 48 maps per group, the intact group used recordings from before the session’s first nonzero stimulation pulse onset marker. The six retained evaluation windows covered [58, 64) seconds after the start of warm-up. The unstimulated group used measured recordings from physical trials 15 and 18. Neither of the two trials had any nonzero stimulation output. From each trial, the group used onesecond windows starting 2, 7, and 12 seconds after the silence event. The two trials had been preceded by 30 and 36 nonzero stimulation pulse onset markers, respectively. The device markers provided event counts. The time of tissue attachment remains undocumented. This comparison cannot separately estimate the effects of signal content, acquisition period, prior stimulation output, and electrode–tissue coupling.

### S3.5 From decision batches to policy and value network updates

With the MEA connected, the policy and value networks shared the current neural readout. The policy network used it to select an action. The value network used it to estimate how many more points could be earned from the current state. Figure 2 shows both inputs. If the readout were masked at only one input, the other could still use the signal, and the between-group comparison could not answer the original question.

**Figure 2.**
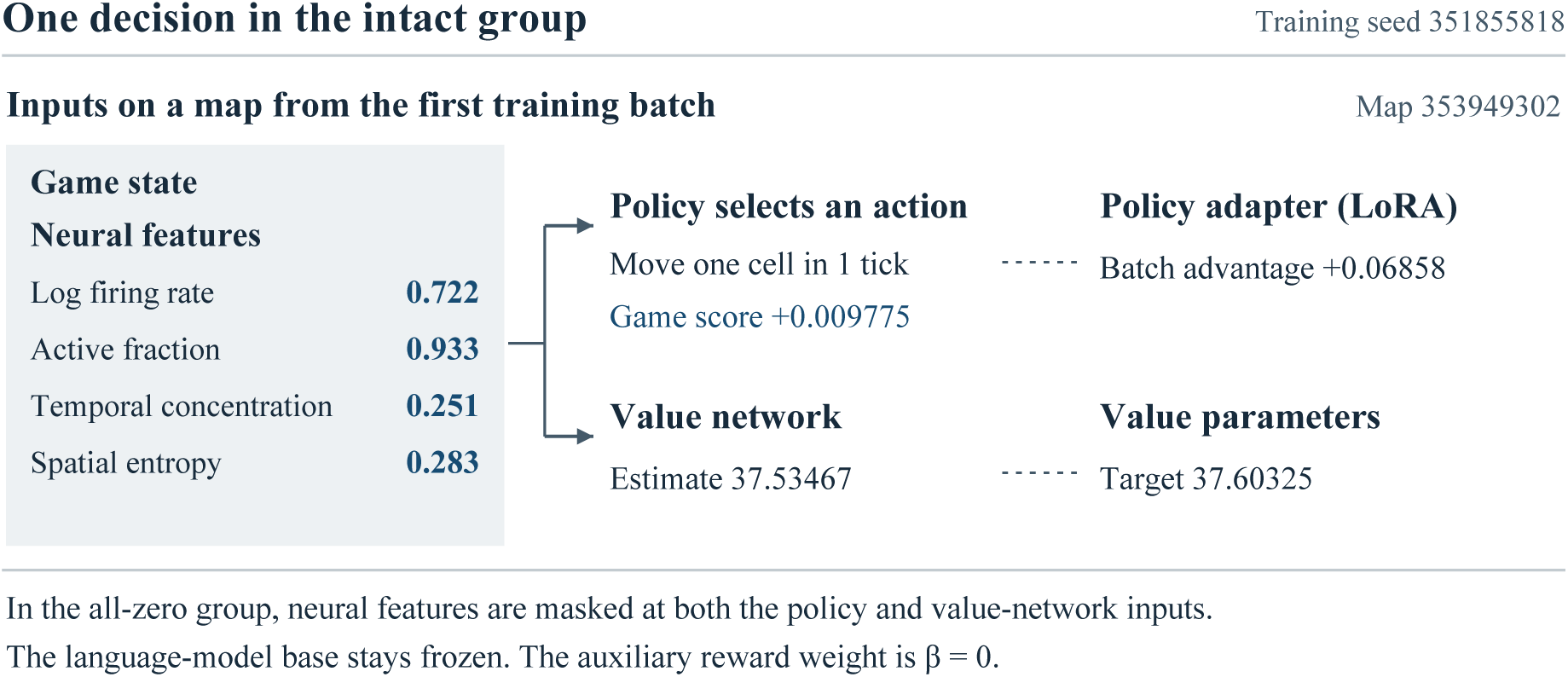
Decision 1 on training map 353949302 in the first batch, for training seed 351855818 in the intact group of the adverse-start study. The policy and value networks received the same set of neural readings. After moving, the character earned 0.009775 game-score points. The advantage estimated from the full batch of trajectories was +0.06858. Adding it to the value estimate for that step yielded a value target of 37.60325. Training updated the policy adapter and the value network while the language-model base remained frozen. This single recorded decision has no confidence interval or between-group test.

#### Algorithm 1

Training batch updates for the adverse-start trial with the second organoid recorded on chip 22021

01 FOR each training seed and input group, the shared initial parameters were loaded.

02 FOR each batch (4 batches total), the game was reset to the registered training map and 12 decisions were collected.

03 Select four neural readings according to the input group and feed them, together with the game state, into the policy and value networks.

04 Sample an action and a target, and record the actual elapsed time d, game score r, value estimate V, and termination flag.

05 The batch was traversed backward. On death, the successor value V*^′^* was set to 0. If the agent was still alive at the end of the batch, the successor value was estimated from the successor state.

06 The temporal difference was *δ* = r + *γ^d^*V*^′^ −* V. At the end of the batch or on termination, A = *δ*. At all other steps, A = *δ* + (*γλ*)*^d^* A*^′^*, where A*^′^* was the advantage at the next step. This duration-adjusted recurrence uses generalized advantage estimation [13].

07 The trajectory was written out. The policy was updated using the advantage A, and the value network was fit to the target V + A.

08 After each batch update, the cumulative step counts of the two AdamW optimizers [14] were checked. The policy count was expected to be 2 times the number of completed batches, and the value count 16 times that number. The final model and optimizer states were saved after the check for batch 4.

The actual logs illustrate how the algorithm worked. In the intact group run with seed 351855818, the agent moved one cell at step 1 of the first batch. The move took 1 game time step and earned 0.009775 points. The value network’s initial estimate was 37.53467. The advantage calculated from the subsequent actions across the entire batch was +0.06858, and the corresponding fitting target was 37.60325. In the equation, *γ* = 0.99985 and *λ* = 0.999. Both values were adjusted for game time steps.

### S3.6 Comparing frozen models on the same maps

The adverse-start trial using recordings from the second organoid recorded on chip 22021 was checked against the available written and machine registration drafts. Each training seed corresponded to three models, one for the intact group, one for the all-zero group, and one for the shuffled group with two-second blocks reordered within trials. During evaluation, model parameters were frozen, each group retained its own input scheme, and runs began in the same scenarios and on the same maps. Figure 3 compares the two evaluation budgets. Section S8 describes the differences between the drafts and the execution labels, as well as the scope of the records that could be independently verified before unblinding.

**Figure 3.**
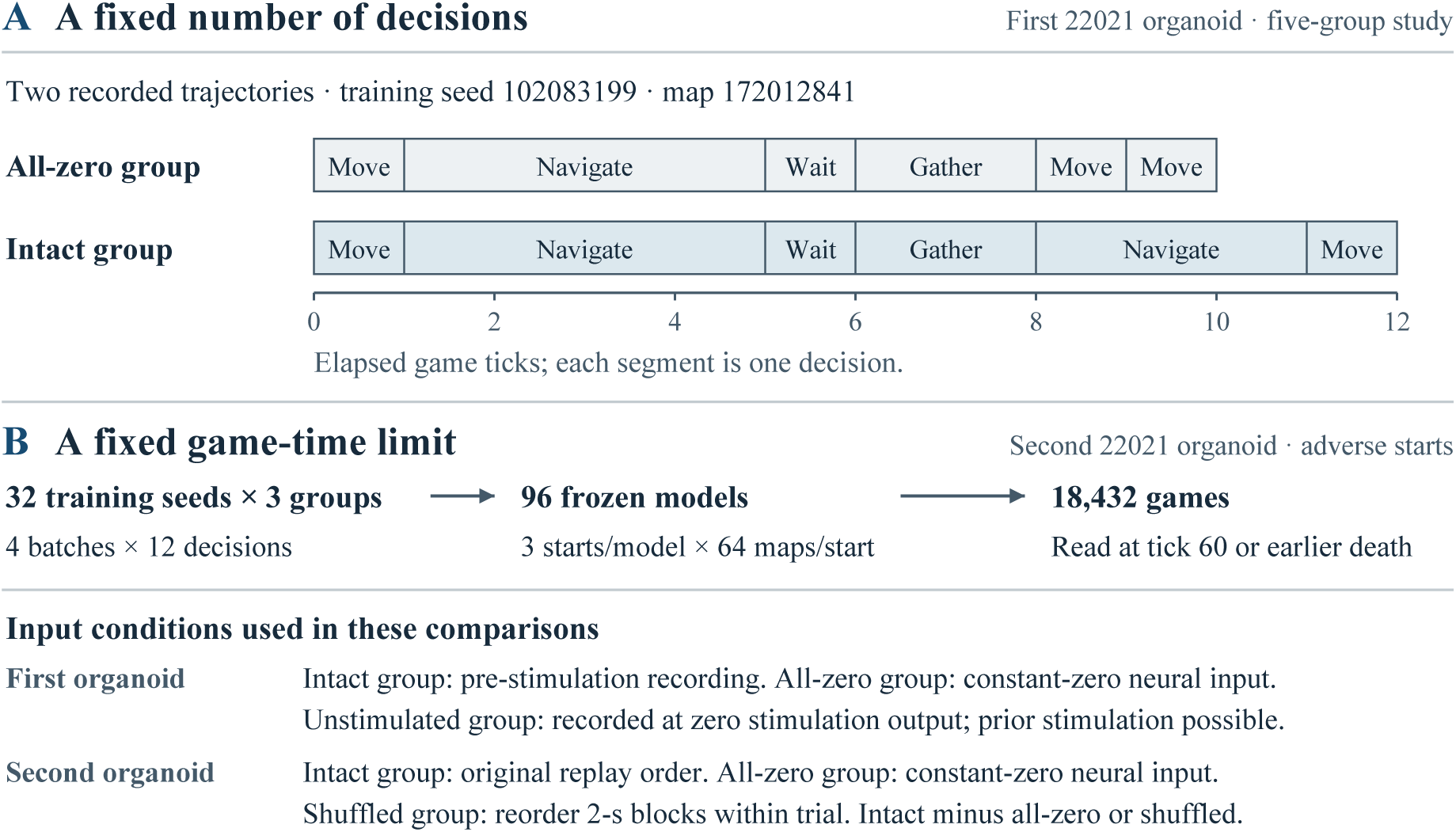
The two experimental rounds evaluated exploration under their respective budgets. (A) Under a fixed decision budget, the panel shows two trajectories from the five-group study of the first organoid on 11/18, recorded during evaluation with training seed 102083199 from the first batch on map 172012841. Each segment represents one decision, and its width shows the duration in game ticks. The all-zero group received constant zero neural input, and the intact group received a recording of spontaneous activity made before stimulation. The groups chose identical actions and targets for the first four decisions. At the fifth decision, the all-zero group chose a single-cell move that ended at game tick 9, while the intact group chose landmark navigation that ended at game tick 11. In the first-organoid study, the unstimulated group received activity recorded while stimulation output was zero. Prior stimulation was possible. (B) The adverse-start study of the second organoid on chip 22021 used a fixed game-time limit. The intact group replayed the input in its registered original order, the all-zero group received constant zero neural input, and the shuffled group reordered two-second blocks within each trial. For matched starts, training seeds, and maps, the differences were calculated separately as intact minus all-zero and intact minus shuffled. The adverse-start endpoint was read at tick 60 or at earlier death. The example trajectories and study diagram show no confidence interval or between-group test.

#### Algorithm 2

Calculation of paired differences in adverse-start endpoints on held-out maps

01 FOR each scenario, training seed, and held-out map, the respective frozen models for all three groups were retrieved.

02 All groups start from the same game state. The intact group replays the held-out windows in the registered order. The shuffled group uses the permutation specified for that evaluation map to reorder the two-second blocks within trials and replays the reordered blocks. Set the all-zero group’s model neural inputs to zero.

03 Actions were executed step by step, and readings were taken at game tick 60 or on death, whichever came first. An action that extended beyond game tick 60 was truncated using the trajectory.

04 The newly revealed cells in the episode were tallied, with each cell counted only one time. The outcome and the drinking and eating events were saved at the same time.

05 In each scenario, paired differences were calculated separately for comparisons with the all-zero group and the shuffled group. For exploration and survival, the difference was the intact group minus the control group. For restricted waiting time, the difference was the control group minus the intact group.

06 Perform crossed resampling of the seeds and shared maps [11] 40,000 times, using the same resampled seed and map indices for all groups in each resample. For each endpoint and seed, calculate the mean paired difference across the held-out maps in the direction specified in the preceding step. Count seeds with a mean paired difference strictly greater than zero as favorable. For each item, report the mean paired difference, the adjusted interval, and the number of favorable seeds.

Each of the 96 frozen models was evaluated under 3 starting conditions, on 64 maps per condition, for a total of 18,432 games. For the low-health start, the primary endpoint was the number of newly revealed cells at game tick 60 or at death if death occurred earlier. For the severe-thirst and severe-hunger starts, the primary endpoint was the survival rate at game tick 60 in each case. The six primary comparisons subtracted the all-zero group and the shuffled group, respectively, from the intact group. Each comparison used a two-sided 99.167% interval. The available registration draft required the lower bound of each interval to exceed zero and at least 75% of seeds to favor the intact group, corresponding to at least 24 of 32 seeds. Overall success required all three endpoints to meet these criteria against each of the two controls. The two secondary endpoints were restricted waiting time to first successful rehydration in the severe-thirst start and to first successful feeding in the severe-hunger start. Each endpoint was compared against two controls, for a total of four comparisons. The denominator for each group included all games. A waiting time of 60 game ticks was assigned if death occurred before the first success or if success had still not occurred by the cutoff. Section S5.6 describes how successful events were identified. Differences were computed as the control group minus the intact group. A positive difference indicated a shorter mean restricted waiting time for the intact group. The draft’s secondary-endpoint provision specified two-sided 98.75% intervals, and its general provision on consistent improvement specified the 75% threshold for favorable seeds. For each of the four waiting-time comparisons, the archived analysis program required the lower interval bound to exceed zero and at least 24/32 seeds to favor the intact group. Overall success was still determined by the six primary comparisons above.

### S3.7 Tracing model parameters from training logs

Figure 4 and Algorithm 3 trace the training logs and the final checkpoint for one model. The intact group (ordered) for training seed 351855818 served as an example. The four per-batch logs and the final parameter snapshot (checkpoint) were stored under models/351855818/ordered/. Each batch log recorded the actions taken by the policy at the time. After four batches of updates were completed, the model’s final parameters and optimizer state were saved in the same checkpoint directory.

**Figure 4.**
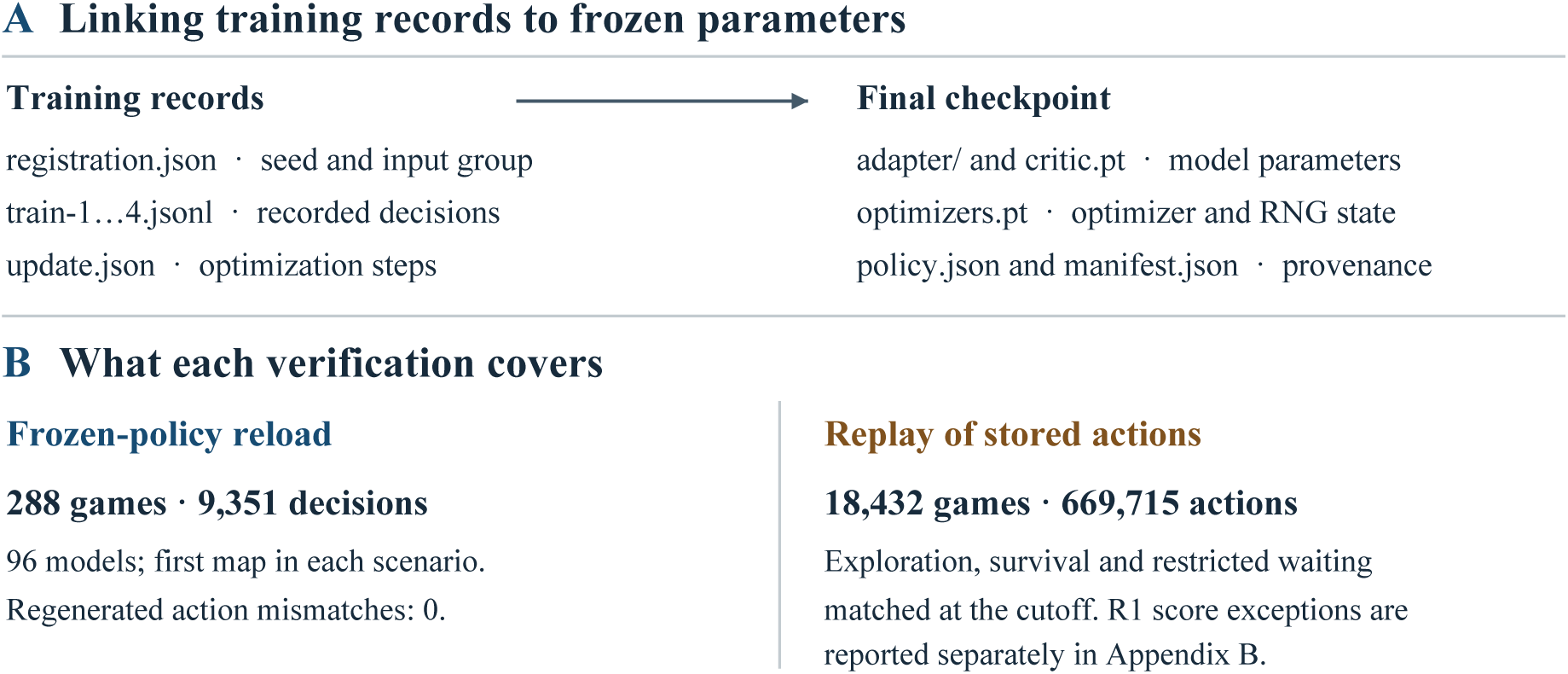
Training logs, final parameters, and replay verification. (A) The training logs train-1.jsonl through train-4.jsonl correspond in order to rounds[0] through rounds[3] in update.json, which records the optimization steps. The registration.json file records the seed and input group, adapter/ and critic.pt contain the model parameters, and optimizers.pt stores the optimizer and random number generator state. The policy.json file and checkpoint manifest record provenance. The checkpoint/manifest.json file stores the training registration fingerprint and the file fingerprint for each parameter component. (B) Frozen-policy reload covered 96 models on the first map of each scenario, totaling 288 games and 9,351 regenerated decisions, with 0 regenerated action mismatches. Replay of stored actions covered 18,432 games and 669,715 actions. In the replay, the exploration, survival, and restricted waiting endpoints matched at the cutoff. Appendix B reports the game-score exceptions separately. These are verification counts. No statistical interval or between-group test applies.

#### Algorithm 3

Verification of the final checkpoint against the batch-by-batch training logs 01 D *→* models/{training seed}/{input group}/, C *→* checkpoint/ under D.

02 reg *→* the contents of D/registration.json, u *→* the contents of D/update.json.

03 FOR b = 1, 2, 3, 4, D/train-{b}.jsonl and the corresponding u.rounds[b *−* 1] were read.

04 The map seed on each log line was checked against world for that batch, and the line count was checked against decisions.

05 m *→* read C/manifest.json. Verify that m.update = 4.

06 Verify that the SHA-256 fingerprint of the canonicalized reg matches m.registration_sha256.

07 Verify the fingerprint of each file in m.files and confirm that u.after.actor = m.actor_sha256.

08 Load the final C/adapter/ and C/critic.pt on the retained evaluation map. Use the frozen model to generate actions and compare them with the stored evaluation actions for that map.

All 12 log lines in the first batch of the training example came from map 353949302, consistent with the update summary for that batch. All 576 component files for the 96 final models passed fingerprint verification. The frozen evaluation was separately spot-checked on map 352055481. All 9,351 actions across three scenarios matched. This spot check covered the final checkpoints. The training trajectories and update processes for the intermediate policies across the four batches still lacked an independent rerun. Appendix B describes the separate full verification by replaying saved actions and the differences in scoring at the cutoff.

## S4 Experimental Setup

### S4.1 Specimens, Chips, and Data Acquisition

A standard MaxOne microelectrode array was used. MaxLab sampled voltage at 20 kHz. The device record showed a spike-detection threshold setting of 5. The units of the threshold field and the conversion to standard deviations remain to be verified. For each specimen, the controller calculated a two-hour operating window from the start time in the software binding record. The time when the tissue was physically connected to the array remains to be verified. Recordings were made from two organoids, one on 11/18 and the other on day 19 of the same month. Both recordings used one chip (22021). A comparison used one additional chip (22554). Chip 28080 was used for single-site pilot training, and the records for chips 28073 and 27701 were used only for initial screening (Table 1).

**Table 1.** Chips and recordings tracked in this study and their uses. Channel counts refer to the mapping, screened set, or candidate set specified in each row. Test episode counts follow the final evaluation criteria.

| Chip / date | Channels and time base | Comparisons included | Availability |
|---|---|---|---|
| 22021 11/18 | There were 322 non-stimulation channels and a 64.99-second baseline. Screening in all three warm-up windows used one common batch of 322 channels. The respective pass counts were 243, 241, and 230 (75.5%, 74.8%, and 71.4%), all below the 95% real-time threshold per window. | The six-decision comparison included 3 × 768 games. The comparison across five groups included 8 × 48 games per arm. | Recordings were available for pre-stimulation spontaneous activity, zero output in that round, and the two stimulation types. |
| 22021 11/19 | There were 324 non-stimulation channels and a 65.00-second baseline. | Each of the five groups had 8 × 48 games. The adverse-start trial had 18,432 games. | The same chip as on the preceding day, with a different organoid. |
| 22554 (a different chip). A scan ID was used on the first day and a temporary session ID on the second day. The number of independent biological samples across the two days remains to be verified. | On 11/19, screening in the original six windows and the two replacement windows used 685 candidate channels for each of H, W, and F. The intersections across the six windows contained 3, 19, and 10 channels, respectively, all below the 50-channel threshold. The intersections across the two replacement windows contained 491, 486, and 474 channels, respectively, and were for that day’s offline comparisons only. Each recording lasted approximately 90 seconds. | Replay from 11/18 was used for the six-decision comparison covering 3 × 768 games and for a recheck of the same batch at game tick 6. The replacement windows from 11/19 were used for the 48-tick and 192-tick offline comparisons. | On 11/18, there were 3 trials per site and 9 trials in total. On 11/19, each site received one stimulation event, and the two time budgets were analyzed separately. |
| 28080 | There were 805 valid mapped hardware channels, counted by hardware channel ID in the original H5 mapping table. | Single-site pilot training with eight training seeds | Offline training and retained evaluations were also archived. Section S4.1 reports the two paired results from the single-site pilot and specifies which data were used for the primary comparisons. |
| 28073 | There were 372 preselected ROI hardware channels for recovery quality control, counted from the ROI channel list in the multisite screening plan. | Preliminary screening | Failed recovery quality control |
| 27701 | After deduplication across wiring configurations, the count was 944 candidate physical electrodes with confirmation recordings. The count was checked by comparing the union of electrode IDs from the initial screening with the list of electrodes covered by the confirmation recordings. | Preliminary screening | 0 candidates met the site screening criteria. |

For the source of the human embryonic stem cell line H9 (WA09), see the ethics statement in Methods. The two-hour period above was the on-site operating window that the controller calculated from the start time in the software binding record. The time when the tissue was physically connected to the array remains to be verified.

Chip 22021 was used to record from two organoids. The recording from the organoid used on 11/18 had a quiet baseline of 64.99 seconds and a map of 732 channels. After the routing was reconstructed, a fixed set of 322 non-stimulation channels was used for response analysis. The organoid was replaced on 11/19, and the recording from the new organoid had a baseline of 65.00 seconds and 324 non-stimulation channels. During baseline, no stimulation was delivered and only spontaneous activity was recorded. Site selection, standardization parameters, and response comparisons were all based on this baseline segment. The old mapping in the service cache differed from the H5 file for that recording, with 1,012 channels in the cache versus 732 in the file. The H5 file took precedence throughout.

Each of the three warm-up check windows for the first 22021 specimen screened the same 322 channels from one batch. Recalculation from the archived raw voltage windows yielded 243, 241, and 230 passing channels, respectively (75.5%, 74.8%, and 71.4%). All were below the original real-time training threshold of 95% per window. Training that day used offline replay after acquisition. The reason for the on-site switch remains to be verified. After acquisition on 11/19, 22554 underwent screening across six windows. Candidates were identified from the recording labeled “silent” in the archive, with 685 candidate channels each at H, W, and F. Each site required at least 50 channels to pass all six windows. The intersections contained 3, 19, and 10 channels, respectively, all below the requirement. The original screening used a one-second reference window beginning 12 seconds before stimulation. It required a median absolute deviation (MAD) ratio relative to the reference window of 0.67 to 1.50 and an absolute median shift of at most 5 ADC counts. All bounds were inclusive. The retained recalculation program required the reference window MAD to be greater than zero. A channel was excluded if any ADC sample in the reference or check window equaled the lower or upper limit of the ADC range. The version of the module loaded on the workstation at the time remains to be verified. With the onset marker for nonzero pulses as time zero, the six one-second windows began at *−*10, *−*7, *−*4, +12, +32, and +52 seconds, respectively. The per-window pass counts were 15, 13, 7, 14, 12, and 19 for H, 127, 71, 69, 75, 77, and 71 for W, and 115, 73, 72, 74, 67, and 93 for F. The alternative screening used the same candidate set at each site, with the earlier window [*−*3, *−*2) seconds as the reference. It took the intersection of the channels that passed in that window and in the later window [+12, +13) seconds. When the earlier window was compared with itself, 685/685 channels passed at each of the three sites. The intersection therefore matched the set of channels that passed in the later window at each site, with 491, 486, and 474 channels, respectively. For each of the two screening procedures, the archived recalculation documented the per-window pass counts and listed the channels in each site’s intersection. The alternative sets were used only for the offline comparisons at game ticks 48 and 192 that day. The six-decision comparison for 22554 and its recheck at game tick 6 used the replay records from 11/18.

In the eight-seed single-site pilot for 28080, the post-stimulation minus pre-stimulation R1 score difference was *−*0.006883 points (95% interval [*−*0.074936, 0.059320]). The post-stimulation minus no-signal difference was *−*0.009394 points ([*−*0.098805, 0.079071]). Each of the two intervals crossed zero, so the score effects were inconclusive. In a separate acquisition session on 11/18, 28080 fell short of the real-time training threshold for that session. The recording from that session was subsequently used to complete offline game training and the retained evaluation. The design and inputs for the three-group low-health start protocol on 11/19 were archived. Whether the evaluation was completed remains to be verified. The formal six-decision comparison involved 22021 and 22554. The groups included in the comparison and the summary results for the two chips are listed in the groups field of summary.json. Per-model training records are in per_model.csv, cited in Section S4.3, and can be located by trial, group, and training seed using the study, arm, and seed fields, respectively. Per-map evaluation records were matched by chip, group, and training seed using the chip, arm, and train_seed fields, respectively. The machine-readable draft registration and the design for carrying out the evaluation of the adverse starts each had one design.json file. Each of the two files identified 22021 as the source of the recordings. The two R1 results for 28080 described above are listed separately in the manuscript’s data summary table under “28080 single-site independent training.” When the scope of data extraction was defined, what criteria were used, and why that scope was chosen remain to be verified.

### S4.2 Stimulation protocol and sites

The original plan was to set the amplitude for each electrode to P90 of its own discharge amplitudes multiplied by 1.2. The calculated amplitudes for the candidate electrodes on 22554 ranged from 45 to 52 µV, below the device’s minimum nonzero setting of 2,982 µV. On each of the two days, the run logs for 22021 listed the electrode coating as platinum black. The material certificate for the chip lot still needs to be verified. Table 3 on page 2 of the manufacturer’s guide [10] lists the recommended starting parameters for standard MaxOne/6WP platinum-black electrodes as 300 to 600 mV, with a phase width of 200 µs and a pulse interval of at least 300 ms. On each of the two days, biphasic commands were sent with the positive phase first, at 199.79 mV and 200 µs per phase. In the regular condition, successive pulse onsets on the same electrode were 200 ms apart, based on the device commands and event markers. The basis for converting the guide’s amplitudes to per-phase DAC commands still needs to be verified. The actual waveform and peak amplitude under electrode load were not measured on site, and actual tissue exposure still cannot be determined.

The original plan was to select the site based on the lowest of health (H), hydration (W), and satiety (F), and to represent the deficit with 0, 1, or 3 pulses. In practice, the five sets of trials used fixed physical conditions. In the regular condition, three pulses were sent to electrode 9393, with onsets at 0, 200, and 400 ms. In the random condition, each trial delivered one pulse at each of three sites. Both organoids were recorded on chip 22021. The 11/18 acquisition session for the first organoid used sites 9393, 2054, and 10783, and the 11/19 acquisition session for the second organoid used sites 9393, 4498, and 2679. Both the site order and the timing of the middle pulse varied. The zero-output condition used the same routing, recording duration, and event markers, with no non-zero pulses sent on those trials. Each of the six permutations of the three conditions was run one time, for a total of 18 segments per day. Baseline screening listed 234 candidate electrodes on 11/18 and 231 on day 19 of the same month. In the actual site order given above for each day, the downloaded routing mapped the sites to hardware channels 49, 676, and 64 on the first day and 49, 956, and 516 on the second day. Replay features were taken from fixed sets of 322 and 324 readout channels that were not used for stimulation, respectively. Conditions were assigned to the same trial numbers on the two days. The regular condition used trials 1, 6, 9, 11, 13, and 17. The random condition used trials 3, 5, 7, 12, 14, and 16. The zero-output condition used trials 2, 4, 8, 10, 15, and 18. The first four blocks were used to extract training windows. The condition orders were regular/zero-output/random, zero-output/random/regular, random/zero-output/regular, and zero-output/regular/random. The last two blocks were used to extract heldout windows. Their condition orders were regular/random/zero-output and random/regular/zero-output, respectively. Each of the six permutations occurred one time in each day’s full acquisition sequence. In chronological order within each condition, the cumulative counts of nonzero pulse-onset markers before the two held-out trials were 24 and 33 for the regular condition, 27 and 30 for the random condition, and 30 and 36 for the zero-output condition. All were device event counts. The prior archival verification cross-checked the H5 files, downloaded routing, transmission acknowledgments, and event markers for each trial on each of the two days. It covered recording segments, site order, relative device frames, and silence markers. The accompanying trial-by-trial details and source fingerprint list provide the item-by-item matches and the fingerprints of the original files, respectively. The waveform and peak amplitude under load at the stimulation electrodes were not measured, and actual tissue exposure still needs to be verified. The regular and random conditions differed in both spatial location and timing.

On 11/18, the validation receipt for the fixed readout panel of 22554 recorded a validation failure. For the exploratory acquisition that followed, the startup check confirmed the panel validation result before the first trial, and the acquisition was logged as not admitted in the acquisition protocol’s real-time admission field. Each block contained one trial for each of H, W, and F, with 3 trials per site in total. Only after acquisition ended was exploratory leave-one-block-out classification performed separately on the pre- and post-stimulation recordings, pooling all 9 trials. Both pre- and post-stimulation classification yielded 5/9 correct predictions, with 1/3, 3/3, and 1/3 correct for H, W, and F, respectively. The exact permutation results were p=0.25 before stimulation and p=0.30556 (approximately 0.306) after stimulation. The reference criteria for the qualification procedure were at least 7/9 correct after stimulation, at least 2/3 correct at each site, post-stimulation p*≤*0.05, and pre-stimulation background p>0.05. The post-stimulation result did not meet these criteria, and the qualification procedure was not invoked for this exploratory acquisition. For game replay, the 6 trials from the first two blocks were used for training, and the 3 trials from the third block were held out.

The pre- and post-stimulation classification results came from the exploratory_classification.pre and exploratory_classification.post fields of the replay archive, respectively. For game replay, the training and held-out block assignments came from the study.train_blocks and study.test_blocks fields of the same archive, respectively.

Hardware verification covered DAC commands, ADC readback, and the sites and timing of stimulation artifacts. Waveforms under electrode load were not measured on site. Actual tissue exposure remained to be verified.

### S4.3 Scale and Randomness

The six-decision comparison on two chips used 16 training seeds per arm, each evaluated on the same 48 held-out maps. The two subtrials with five groups each were conducted on two separate days, with 8 seeds per arm per day and the same set of 48 maps. The adverse-start trial on the second organoid recorded on chip 22021 used 32 seeds per arm. Each of the three start conditions was evaluated on 64 shared maps. The training and evaluation maps for this trial did not overlap, and the groups shared the evaluation maps and action-sampling seeds. Evaluation with fixed parameters and subtraction within matched seed–map pairs can control for shared map conditions. The differences still varied by map. All seeds were algorithmic replicates.

The interaction budget for the six-decision design was 4 batches of 12 decisions each. The per-model registry for the 48 models across two cohorts and three groups on the first specimen of 22021 showed agreement on all four initial weight fingerprints, the game and reward configurations, and the optimization hyperparameters. The update receipts for all 192 batches recorded 2 policy update steps, no KL early stopping, and zero auxiliary reward credited. The final receipt for each model recorded 8 policy optimizer steps and 64 value optimizer steps. The weight fingerprints in the receipts were limited to registry entries. Runtime fingerprints for the actual processes’ entry points, dependencies, devices, and numerical precision were still missing. According to the accompanying per-model training budget manifest (per_model.csv), the initial and expansion cohorts in each of the two six-decision trials each contained 8 models per group. Each model completed 4 collection batches of 12 decisions each, for a total of 48 training decisions, 8 policy optimization steps, and 64 value optimization steps. The initial and expansion cohorts were summarized separately by group for each trial. Within each cohort, the 8 models in each group completed a combined 384 training decisions, 64 policy optimization steps, and 512 value optimization steps. No batch update receipt in any group recorded KL early stopping. For 22021, the training game tick counts for the intact, all-zero, and unstimulated groups were 567, 538, and 561, respectively, in the initial cohort, totaling 1,666 game ticks. In the expansion cohort, the respective counts were 564, 563, and 571 game ticks, totaling 1,698 game ticks. For 22554, the training game tick counts for the intact, shuffled, and native R1-only groups were 582, 582, and 574, respectively, in the initial cohort, totaling 1,738 game ticks. In the expansion cohort, the respective counts were 557, 550, and 534 game ticks, totaling 1,641 game ticks. The per-batch value update counts for the first specimen of 22021 were supported by checks of cumulative counts at the end of each batch and by the final state. The update receipts for 22554 also included a field for the per-batch value update count. The manifest used the trial, group, and training seed (study, arm, seed) to locate per-model training trajectories and update receipts, then linked them to per-map evaluation records by chip, group, and training seed (chip, arm, train_seed). Independent reproduction of the per-batch training updates, complete runtime fingerprints of the successful processes, elapsed training time on the original machine, and GPU-hours remained to be verified.

The adverse-start experiment on the second specimen of 22021 included an additional 96 models. The 384 archived batch trajectories contained a total of 4,608 training decisions and 6,681 game ticks. The all-zero group, intact group, and shuffled group each had 32 models, with 128 batches and 1,536 training decisions retained per group. The three groups recorded 2,179, 2,253, and 2,249 training game ticks, respectively. The registered configuration specified 2 policy training epochs and 16 value training epochs per batch and a minibatch size of 12 for both networks. The policy accumulated gradients within each minibatch using microbatches of size 4. The per-batch update receipts recorded 2 policy update steps per batch, with no KL early stopping in any batch. Checks of the cumulative AdamW step count at the end of each batch and the final snapshots supported the count of 16 value update steps per batch (see Algorithm 1). Each model accumulated 8 policy update steps and 64 value update steps. Each group accumulated 256 policy update steps and 2,048 value update steps. Archive verification covered the actual training trajectories, update receipts, and final states. Independent reproduction of the per-batch updates remained to be verified. See Appendix B for spot checks of the frozen-model evaluations.

In the first-day experiment with five groups, each group had 8 training seeds, and each seed was evaluated on 48 maps. The intact group used the spontaneous activity recorded before the first stimulation. The unstimulated group used the window actually recorded when there was no nonzero output in that round. The specific window timings listed in Section S3.4 applied to the first specimen’s evaluation with three groups and six decisions. According to the input re-extraction verification summary, verification for the adverse-start experiment on the second specimen covered 24 calibration windows, 48 training windows, 4 windows for end-of-batch estimates of subsequent value, and 60 retained evaluation windows. That experiment used 64 evaluation maps per seed per scenario. Recording windows and maps were counted separately. Source windows and model logs were also checked separately for the five-group experiment on each of the two days. The intact group’s batches started 2, 15, 28, and 41 seconds after warm-up, in that order. Starting at each of those times, 12 consecutive one-second training windows were taken, and the next window was reserved for the end-of-batch value estimate. The retained windows covered the interval [58, 64) seconds after warm-up. In each of the first four physical blocks, the regular, random, and unstimulated groups used the trial for their respective condition and followed the same window schedule starting 2 seconds after that trial’s first event marker. In each of the last two physical blocks, they took one-second windows starting 2, 7, and 12 seconds after the marker. The unstimulated group used the silence marker as its reference point. The all-zero group used the unstimulated group’s window indices with its inputs set to zero. For a given day and group, the same set of retained windows was reused in decision order on each retained map. The training snapshots and retained logs allowed the window indices and inputs to be checked step by step. The designated window and resulting scalar were retained for the end-of-batch value calculation. A separate input snapshot for that call was unavailable.

### S4.4 Statistical methods

For the archived six-decision intervals, the mean paired difference was first calculated for each training seed over a fixed set of 48 maps. Training seeds were then resampled with replacement 12,000 times to obtain two-sided 95% percentile intervals. For the two primary six-decision comparisons on the first specimen of 22021, the archived intervals used the original fixed set of 48 maps and resampled from the 16 seeds pooled across the initial and expansion batches. The proportions of the two batches could vary across resamples. The post hoc sensitivity analysis held fixed the original 48 maps and the allocation of 8 seeds per batch. It sampled 8 seeds with replacement within each of the initial and expansion batches, then pooled the seed-level mean differences from the two batches with equal weights. Exact convolution of the integer-valued distribution of total paired differences in counts of newly revealed cells yielded two-sided 95% percentile intervals. The same batch of trajectories was truncated post hoc at game tick 6. The archived intervals were computed using 20,000 iterations of crossed resampling of seeds and common maps. A 95% interval was reported for each comparison. The sensitivity analysis for *β*=0 also used the original *β* sweep’s nominal Bonferroni level of 98.333% for the three nonzero *β* comparisons. For the primary and secondary endpoints in the adversestart experiment, crossed resampling was performed 40,000 times according to the comparison families in Section S3.6. All arms used the same seed and map indices in each iteration. Intervals based on resampling seeds alone were conditional on the maps tested. Intervals based on crossed resampling of seeds and common maps reflected variation from changing both training seeds and maps. All arms of the adverse-start experiment shared the map indices in each resample. For comparisons involving the shuffled group, each map was resampled together with its assigned replay permutation. Repeated draws of a map retained the same permutation. The resulting intervals applied to the tested maps under the scheme that assigned their fixed permutations. Variation from assigning a different permutation to the same map had yet to be estimated. All intervals were conditional on the biological recordings used in the corresponding experiment.

In the two-day, five-group trial, counts of newly revealed cells and total R1 were analyzed separately by day. Each comparison comprised 384 pairs matched by training seed and map. For each comparison family on each day, we performed 30,000 resampling draws. In each draw, we sampled 8 training seeds and 48 shared maps with replacement and used the Cartesian product of the two sets of indices to compute the mean paired difference. All comparisons and the two endpoints within a family used the same indices in each draw. The primary family for each day comprised the unstimulated group minus the all-zero group, the post-regularstimulation group minus the unstimulated group, and the post-random-stimulation group minus the unstimulated group. Each group pair had two endpoints, the count of newly revealed cells and total R1, for six comparisons in total. We used a two-sided 99.167% percentile interval for each comparison and a resampling random seed of 632201. The supplementary family comprised the two endpoints for the intact group minus the all-zero group. We used a two-sided 97.5% percentile interval for each endpoint and a resampling random seed of 632202. For each training seed, we first computed the mean paired difference across the 48 common maps for each of the two endpoints. A seed counted as favorable on both endpoints only if the two differences each strictly exceeded the numerical tolerance of 10*^−^*^10^. A between-group comparison met the criterion only if the lower bounds of its two adjusted intervals were each above zero and at least 6/8 seeds were favorable on both endpoints. Interval inference was conditional on each day’s biological recordings and the map generation mechanism used. The “Archived Computation and Current Recalculation” section of the accompanying “Interval Resampling Units for the two-Day, five-Group Trial” lists by date the locations of the first day’s offline stimulation control archive and the second day’s offline five-group archive. The primary design for each day is in design.json in the archive root. The supplementary design for the intact group is in design.json in the pre-stim-baseline subdirectory of that same root. In each of the two design files, the comparison families and resampling settings can be checked using contrasts, endpoints, interval.draws, interval.seed, and interval.quantiles. In each day’s compare.py, lines 114–124 specify the pairing and crossed resampling with shared indices, and lines 128–131 specify how seeds favorable on both endpoints were counted. Each day’s comparison.json stores the archived interval endpoints in the interval field indexed by group pair and endpoint. The accompanying index identifies the receipts for the interval recalculation and the independent numerical verification. Across the two days, all 32 endpoints of the 16 intervals for counts of newly revealed cells and total R1 matched the archive. The two time-subtracted residuals for the first day, reported in the final paragraph of Section S5.2, were calculated separately by lines 151–162 of analyze.py in the statistical recalculation supplement. We used random seed 220211119 to perform 30,000 crossed resampling draws of training seeds and shared maps and obtained a two-sided 97.5% percentile interval for each residual. The historical analysis process, NumPy environment, and runtime state of the random number generator remain to be verified.

## S5 Results

### S5.1 Map coverage under the same decision budget

For the first specimen on chip 22021, we compared the count of newly revealed cells after six decisions per episode across 16 seeds and 48 maps. The all-zero group, intact group, and unstimulated group each had 768 episodes. Their totals were 31,017, 32,543, and 32,674 newly revealed cells, respectively, and their means were 40.387, 42.374, and 42.544 cells per episode, respectively. Relative to the all-zero group, the intact group and the unstimulated group revealed 1.99 and 2.16 more cells per episode on average, respectively. The mean difference was positive for 14/16 and 15/16 seeds, respectively (Table 2, Figure 5). In the initial and expanded cohorts, respectively, the mean differences for the intact group minus the all-zero group were +1.818 and +2.156 cells per episode. The corresponding differences for the unstimulated group minus the all-zero group were +2.438 and +1.878 cells per episode. Each comparison had 384 pairs per cohort. The two point estimates were positive in both of the two cohorts. The unstimulated group had the larger mean difference in the initial cohort, and the intact group had the larger mean difference in the expanded cohort. The two mean differences corresponded to 4.92% and 5.34% of the mean count of newly revealed cells in the all-zero group, respectively. Using the sample standard deviation of all episode-level paired differences in each comparison as the denominator, the paired effect sizes dz were 0.196 and 0.206, respectively. Survival was 768/768 episodes in each of the three groups. On the 48 shared maps, the mean difference for the direct comparison of the intact group minus the unstimulated group was *−*0.17 cells per episode. The descriptive mean differences were *−*0.62 and +0.28 cells per episode in the initial and expanded cohorts, respectively, with 384 pairs per cohort. The post hoc two-sided 95% seed-cluster t interval was computed conditional on the fixed map set. The 16 seeds from the two cohorts, with no overlap between cohorts, were treated as seed clusters that could be pooled. The interval was [*−*1.00, 0.66] cells per episode and included zero. The mean differences and intervals for the two primary comparisons at six decisions described the paired results from the runs actually conducted in these two cohorts on the shared maps. It remains to be verified whether the expanded cohort size, stopping rule, and pooled reporting rule were set before the initial cohort results were first examined, and whether those results influenced these decisions. The post hoc analysis held the original 48 maps fixed, resampled 8 seeds within each of the two cohorts, and pooled the cohorts with equal weights. The two-sided 95% intervals for the intact group minus the all-zero group and the unstimulated group minus the all-zero group were [1.204, 2.742] and [1.590, 2.707] cells per episode, respectively. Both intervals were entirely above zero. Inference was limited to this record, the original 48 maps, and a composition in which each of the two cohorts contributed half. The retained results for the 384 episodes per arm in the expanded cohort were all traceable to their respective model registry entries. The windows actually read or the all-zero masks matched the assigned groups. These mean differences described the full set of group conditions as implemented. The receipts for individual models were used to verify the registered configurations and actual update counts. Runtime fingerprints for the code and environment that were actually loaded were still missing. The archived results could not determine why the ordering reversed between the two cohorts.

**Table 2.** Paired results for chip 22021. The first two rows report exploration after six decisions for the first specimen, measured in newly revealed cells per episode, with 768 episodes per arm. The 95% intervals are conditional on the fixed map set. The last six rows are from the second specimen, with 2,048 episodes per arm per scenario. The low-health start results report newly revealed cells. The severe-thirst and severe-hunger start results report the number of episodes with survival and differences in survival rates (percentage points). The last six rows use 99.167% intervals. Within each row, seeds with a positive difference are counted using the same endpoint and control. For adverse starts, all groups share the resampled map indices, and the shuffled group’s fixed replay permutations are resampled together with their maps. Variation from reassigning permutations to the same map has yet to be estimated.

| Starting condition and metric | Experimental group / control group | Observed values for both groups | Paired difference [interval] | Seeds with a positive difference |
|---|---|---|---|---|
| six decisions · exploration | Intact group / all-zero group | 42.374 / 40.387 | +1.99 [1.18, 2.73] | 14/16 |
| six decisions · exploration | Unstimulated group / all-zero group | 42.544 / 40.387 | +2.16 [1.58, 2.75] | 15/16 |
| Low-health start · exploration | Intact group / all-zero group | 131.717 / 126.125 | +5.59 [−0.19, 11.26] | 25/32 |
| Low-health start · exploration | Intact group / shuffled group | 131.717 / 129.438 | +2.28 [−1.92, 7.13] | 21/32 |
| Severe-thirst start · survival | Intact group / all-zero group | 803 / 862 | −2.88 [−9.08, 2.73] | 7/32 |
| Severe-thirst start · survival | Intact group / shuffled group | 803 / 837 | −1.66 [−6.35, 2.83] | 10/32 |
| Severe-hunger start · survival | Intact group / all-zero group | 1,210 / 1,275 | −3.17 [−9.03, 2.34] | 5/32 |
| Severe-hunger start · survival | Intact group / shuffled group | 1,210 / 1,233 | −1.12 [−5.71, 3.29] | 9/32 |

**Figure 5.**
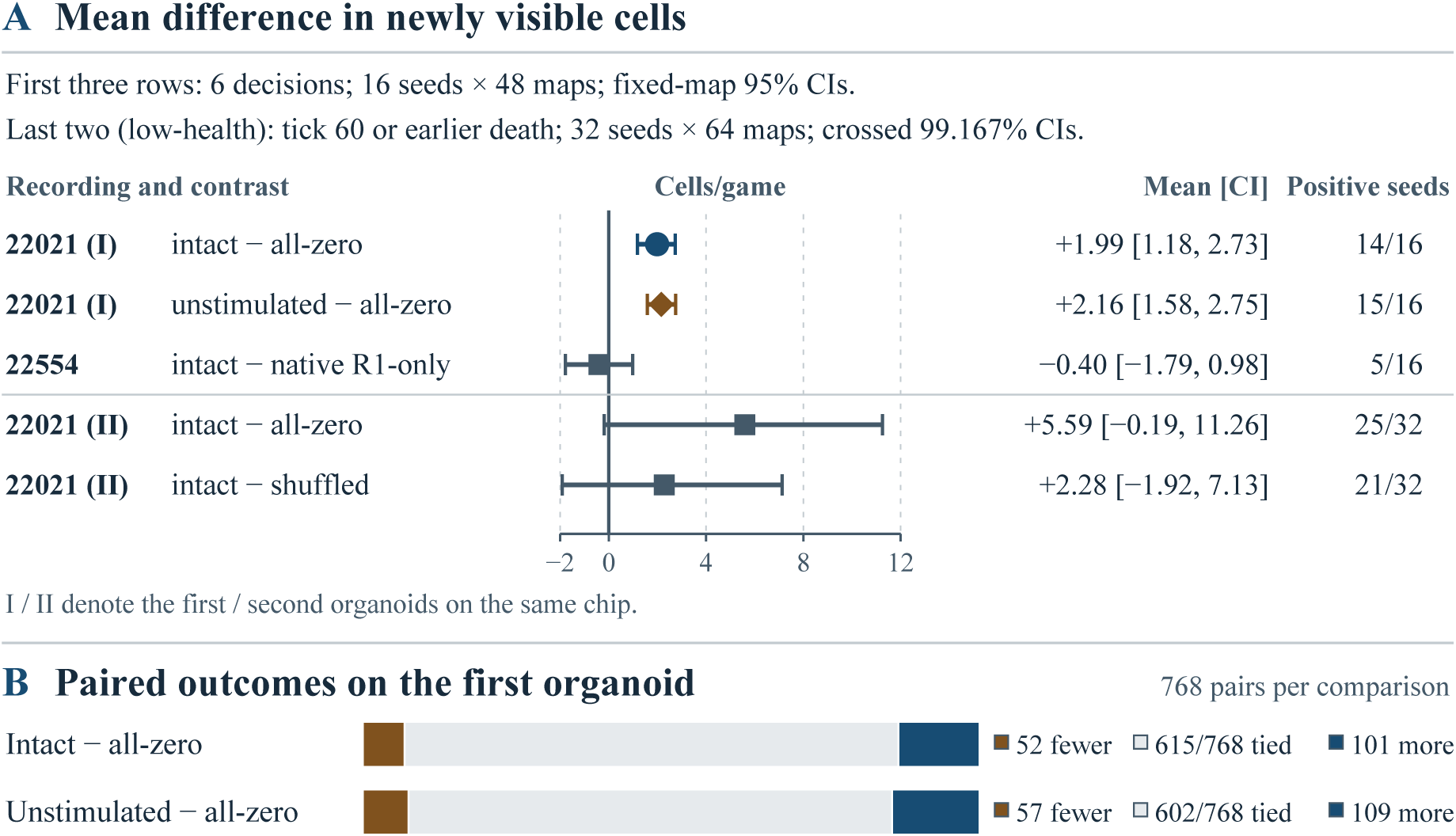
Paired differences in newly revealed cells (cells per game). (A) Points show mean paired differences calculated as the experimental group minus the control group. Horizontal lines show two-sided confidence intervals, the vertical line marks zero, and the right column counts seeds with positive mean paired differences. The first three rows used 6 decisions per game and 16 seeds *×* 48 maps per arm. The 95% confidence intervals were computed by resampling seeds with maps held fixed. The first two rows show the intact group minus the all-zero group and the unstimulated group minus the all-zero group, respectively, for the first organoid on chip 22021. The third row shows the intact group with ordered training pairing minus the native R1-only group on chip 22554. The last two rows show results from the low-health start for the second organoid on chip 22021, with 32 seeds *×* 64 maps per arm. Newly revealed cells were counted at time step 60 or at earlier death. The intact group replayed the original sequence, and the all-zero and shuffled groups were each subtracted from it in separate comparisons. The shuffled group reordered two-second blocks within each trial. The 99.167% confidence intervals used crossed resampling of seeds and maps. (B) For the first organoid, each pair is classified by whether the experimental group had fewer, equal, or more newly revealed cells than the all-zero group. Of the 768 seed–map pairs per comparison, 615 tied for intact minus all-zero and 602 tied for unstimulated minus all-zero. These counts are descriptive.

The intact group and the unstimulated group were each paired with the all-zero group by training seed and map, yielding 768 episode pairs per comparison. In that order, the counts of increases, ties, and decreases in newly revealed cells were 101/615/52 and 109/602/57 for the two comparisons. Using all 768 pairs in each comparison as the denominator, the win, tie, and loss rates were 13.15%, 80.08%, and 6.77% for the first comparison and 14.19%, 78.39%, and 7.42% for the second. The median paired difference was 0 cells in both of the two comparisons. Among pairs with different counts of newly revealed cells, 66.01% of the 153 pairs for the intact group minus the all-zero group had positive differences, as did 65.66% of the 166 pairs for the unstimulated group minus the all-zero group. The cell counts matched across all three groups in 577 pairs. After zero differences were excluded and tied absolute differences were assigned average ranks, the paired rank-biserial correlations were 0.500 and 0.503, respectively. After the largest 38 positive differences were removed from each comparison, the two mean differences shrank to +0.186 and +0.303 cells, respectively (Section S6.2).

### S5.2 Sources of the scores

In the five-group subtrial with the first specimen on 11/18, each of the 8 seeds per group was run on 48 maps. In all 1,920 games, the agent survived to the sixth decision. The number of elapsed game ticks in each game was an integer. The intact group and the unstimulated group were each paired with the all-zero group. Across 384 matched pairs per comparison, the intact group and the unstimulated group lasted 130 and 150 more game ticks in total, respectively, or 0.339 and 0.391 more game ticks per game on average. The survival timer awarded 0.01 points per game tick, with a within-action discount factor of 0.99985 per game tick. The differences in total R1, calculated by subtracting the all-zero group from the intact group and the unstimulated group, were +0.004017 points per game (97.5% interval [0.001292, 0.008037]) and +0.004073 points per game (99.167% interval [0.000866, 0.008420]), respectively. Each of the two differences was positive for 8/8 seeds. The differences in timer points were +0.003385 and +0.003905 points per game, accounting for 84.3% and 95.9% of the corresponding differences in total R1, respectively (Figure 6). Each of these two percentages was based on 384 matched game pairs from the first day.

**Figure 6.**
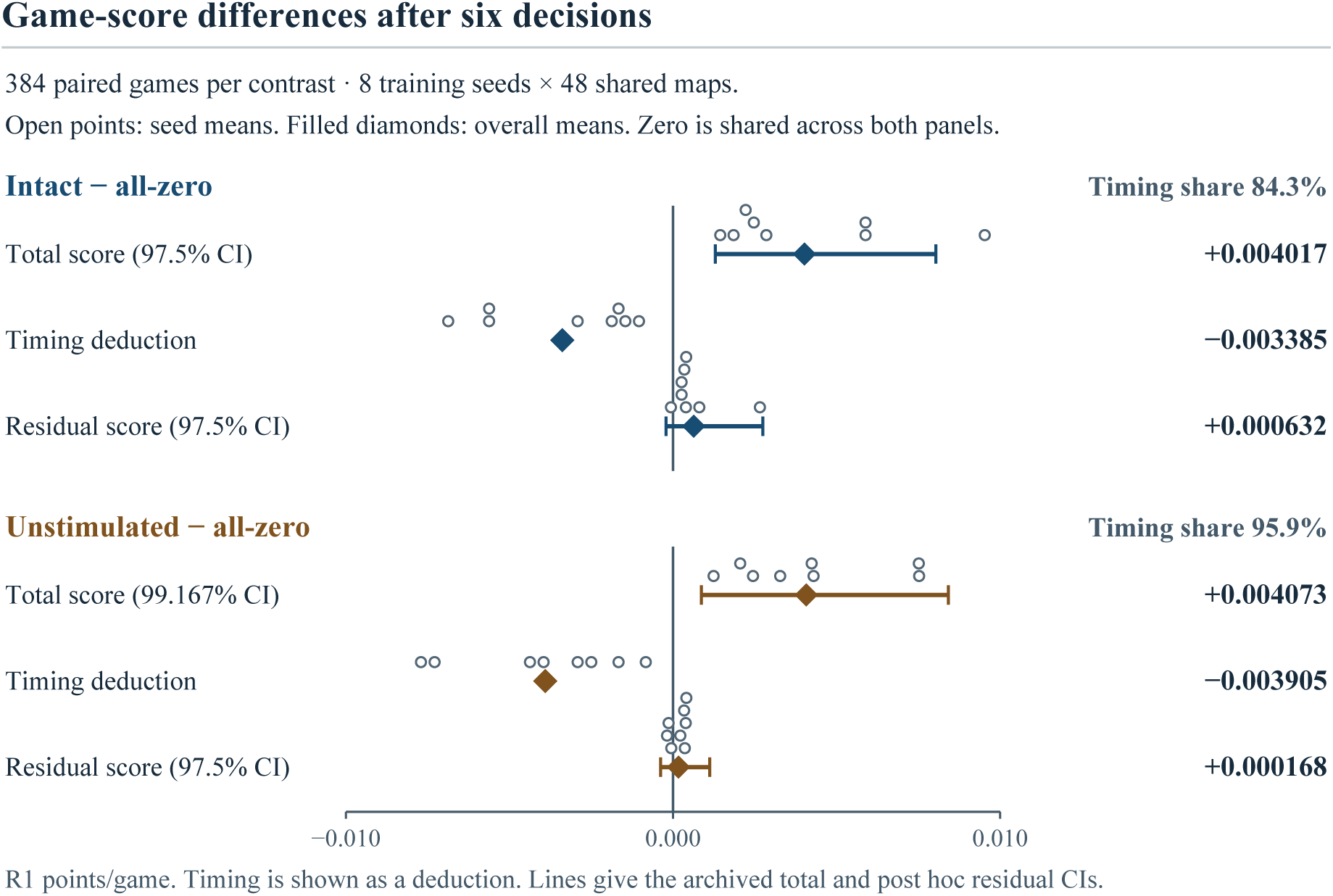
Game-score decomposition for the first sample on 11/18. The first-day five-group experiment is shown through the intact minus all-zero and unstimulated minus all-zero contrasts. The intact group used spontaneous activity before stimulation, the unstimulated group used activity recorded in a trial with zero stimulation output, and the all-zero group had its neural input masked to zero. Each contrast used 8 training seeds *×* 48 shared maps, for 384 paired games per contrast, with 6 decisions per game. The rows show the total R1 difference, the timing difference plotted with its sign reversed, and the residual score difference after subtracting the timing contribution, all in points per game. Each row shows point estimates, with open circles marking seed means and filled diamonds marking overall means. Lines show the archived confidence intervals for the total-score differences and the post hoc confidence intervals for the residual-score differences. For the two total-score contrasts reported in Section S5.2, the archived confidence levels were 97.5% for intact minus all-zero and 99.167% for unstimulated minus all-zero. The total-score difference was positive for 8/8 training seeds for intact minus all-zero and 8/8 for unstimulated minus all-zero. The two post hoc 97.5% confidence intervals for the residual-score differences both crossed zero. The intervals use crossed resampling of seeds and shared maps. No interval is shown for the timing deduction. The timing share is the magnitude of the mean timing deduction divided by the mean total score difference.

After timer points were subtracted from each game’s score, the difference between the intact group and the all-zero group was +0.000632 points (post hoc 97.5% interval [*−*0.000217, +0.002742]), and the difference between the unstimulated group and the all-zero group was +0.000168 points ([*−*0.000382, +0.001124]). The two intervals both crossed zero. The remaining scores still included discounted changes in resources and health status. These scores alone could not be used to assess decision quality.

### S5.3 Changes in action composition

For the first specimen on chip 22021, each group had 4,608 actual action selections. Compared with the allzero group, the unstimulated group selected single-cell movement less often, with the count decreasing from 2,207 to 2,083 (*−*2.69 percentage points). Landmark navigation increased from 394 to 499 (+2.28 percentage points), and gathering increased from 499 to 555 (+1.22 percentage points, calculated from unrounded counts). Table 3 and Figure 7 show the frequencies by action category. These totals alone could not determine how much of the difference in newly revealed cells was attributable to any one action category.

**Table 3.** Observed selection frequencies by action category for the first specimen on chip 22021 under a budget of six decisions. The denominator is 4,608 decisions per arm. The table lists only four categories. No between-group tests were performed.

| Action type | All-zero group | Intact group | Unstimulated group |
|---|---|---|---|
| Single-cell movement | 47.89% | 45.46% | 45.20% |
| Landmark navigation | 8.55% | 10.87% | 10.83% |
| Gathering | 10.83% | 11.89% | 12.04% |
| Waiting | 17.23% | 16.12% | 16.38% |

**Figure 7.**
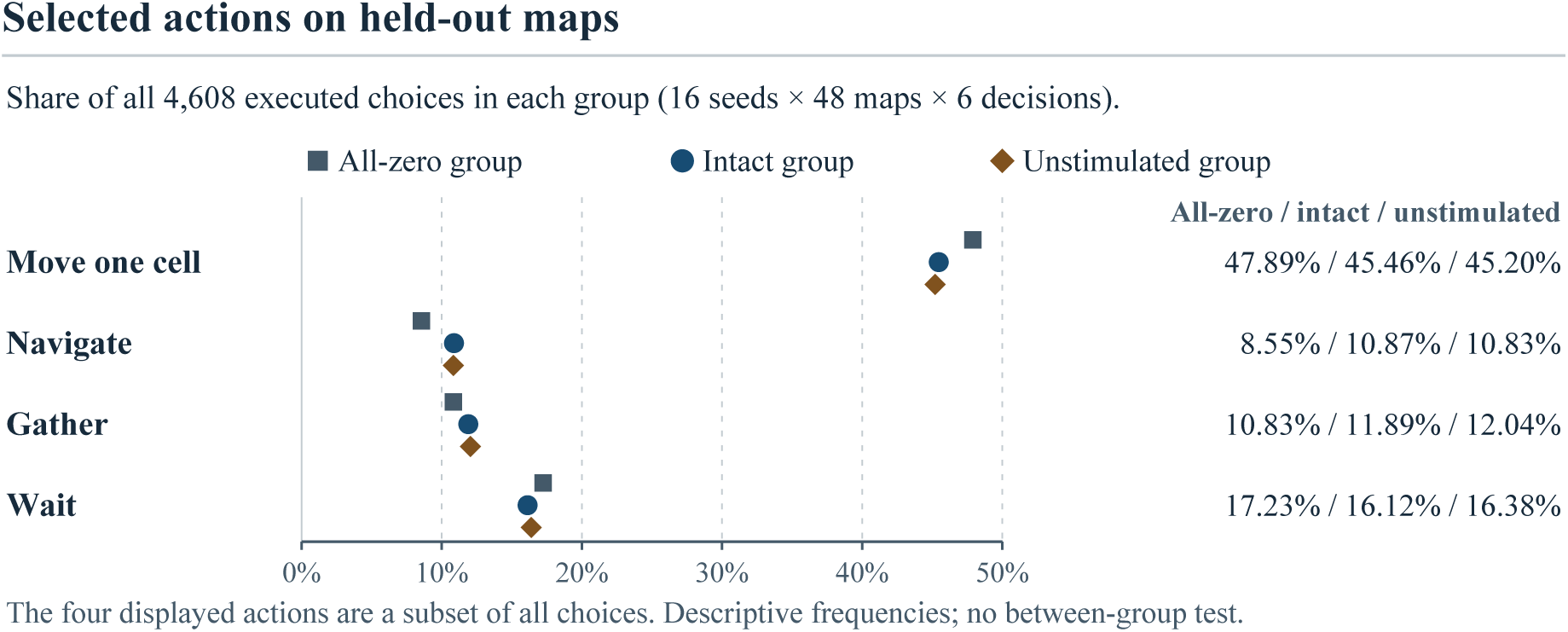
Percentages of all executed choices in each group accounted for by the four displayed action types on the held-out maps for 22021. The legend identifies the all-zero, intact, and unstimulated groups. Each group contributed 4,608 choices (16 seeds *×* 48 maps *×* 6 decisions per seed–map pair). The plotted frequencies are descriptive and use the same choices as Table 3. No between-group tests were performed. Squares, circles, and diamonds distinguish the all-zero, intact, and unstimulated groups, respectively. No interval is shown.

These frequencies use all decisions as the denominator. Determining which cells were actually traversed or how many cells a navigation action newly revealed for the same seed and map would require pairing the visible areas before and after the action. The study did not perform this action-level attribution.

### S5.4 Stimulation comparisons over two days and the second specimen on the same chip

In the two-day, five-group trial, each arm used the same 8 seeds and 48 maps each day. Readouts following the two stimulation conditions, regular and random, were each paired with readouts from the unstimulated group. On 11/18, the differences in newly revealed cells were *−*0.8125 [*−*3.182, 0.906] and +0.1953 [*−*1.495, 1.682] cells, respectively, and the R1 differences were *−*0.000777 [*−*0.003446, 0.001326] and +0.000576 [*−*0.001536, 0.003840] points, respectively. On 11/19, the differences in newly revealed cells were *−*0.4870 [*−*2.760, 1.424] and *−*0.2865 [*−*2.706, 2.401] cells, respectively, and the R1 differences were *−*0.000785 [*−*0.003990, 0.001539] and *−*0.000738 [*−*0.004779, 0.002393] points, respectively. All eight of these 99.167% intervals crossed zero. They described differences in exploration and scores under the data collection and offline replay arrangements actually used on each day. Section S4.2 gives the actual order of the training and retention blocks over the two days. The balance across the six permutations applied to the full acquisition sequence for each day. On both of the two days, all retained windows for the unstimulated group came from the final trials of their respective retention blocks. Each of those trials immediately followed two trials with nonzero stimulation output. Pairing by training seed and map controlled for shared map conditions. The source trials’ positions within blocks and prior stimulation histories still varied by condition. The regular and random conditions also differed in both stimulation sites and timing. These comparisons could not separate the independent effect of stimulation output in the current trial, carryover from prior stimulation, and the contribution of within-session drift.

Under the same six-decision budget in the first day’s five-group experiment on 11/18, the intact group and the unstimulated group were each paired with the all-zero group by training seed and map, with 384 pairs per comparison. The exploration difference for the intact group minus the all-zero group was +1.82 cells per episode (97.5% interval [*−*0.32, 4.61], with 6/8 seeds favoring the intact group). For the unstimulated group minus the all-zero group, the difference was +2.44 cells per episode (99.167% interval [*−*0.16, 6.50], with 8/8 seeds favoring the unstimulated group). The two exploration intervals each crossed zero, and neither comparison met the dual-endpoint criterion for the five-group experiment. Section S5.2 reports the intervals for total R1 and the timing breakdown for the same pairings.

On 11/19, a different organoid was used on the same chip, and a separate acquisition session was started. The second day’s five-group evaluation reused the first day’s 8 training seeds and 48 maps. The initial batch of the pooled six-decision analysis for the first specimen directly included the models and evaluation episodes from the intact group, the all-zero group, and the unstimulated group in the first day’s five-group experiment, with 384 episodes per group. The expansion batch used an additional 8 training seeds and reused the same set of 48 maps. Thus, for each of the first day’s two exploration comparisons, 384 pairs were already included in the corresponding pooled result. Section S5.1 reports the results by batch. The per-model training budget inventory cited in Section S4.3 identifies the model sources. Appendix B describes how the retained episodes were pooled. On the second day, the intact group minus the all-zero group yielded +1.77 cells (97.5% interval [*−*0.66, 5.05], with 7/8 seeds favoring the intact group), and the unstimulated group minus the all-zero group yielded +2.15 cells (99.167% interval [*−*0.72, 6.08], with 8/8 seeds favoring the unstimulated group). The two intervals each crossed zero, so the second-day exploration effects were inconclusive. The corresponding R1 differences were +0.003886 points (97.5% interval [0.001082, 0.007924]) and +0.004412 points (99.167% interval [0.001043, 0.009612]), each with positive differences for 8/8 seeds. These two specimen– session combinations were insufficient to estimate between-specimen variation.

In the order of the unstimulated group minus the all-zero group, the post-regular-stimulation group minus the unstimulated group, the post-random-stimulation group minus the unstimulated group, and the intact group minus the all-zero group, the counts of seeds with favorable differences on both endpoints were 8/8, 1/8, 3/8, and 6/8 on 11/18, and 8/8, 1/8, 3/8, and 7/8 on 11/19, respectively. On each of the two days, all four adjusted exploration intervals crossed zero. For each of the two post-stimulation groups minus the unstimulated group, the adjusted interval for total R1 also crossed zero, and the count of seeds favoring the post-stimulation group on both endpoints fell below the threshold. Thus, on each of the two days, 0/4 comparisons met the full criterion.

### S5.5 Exploration, Score, and Resources for Chip 22554

For 22554, the six-decision evaluation used recordings from 11/18. Each arm used 16 training seeds *×* 48 retained maps, for a total of 768 episodes and 4,608 decision rows per arm. The critic_input_size field read directly from the evaluation logs indicated that the native R1-only group did not append the four-dimensional MEA readings and had a 108-dimensional value network input. The intact group (ordered training pairings) and the shuffled group (shuffled pairings of training trials) each had a 112-dimensional input. The differences obtained by subtracting the native R1-only group from each of the two groups with MEA input encompassed both the addition of the four-dimensional readings and the change in the structure of the value network input. The two groups with MEA input differed in how they paired training trials, and both preserved the withintrial order before and after stimulation. The input fingerprints differed in all 64 pairs of training batches. The direct comparison between the two groups estimated the performance difference, under a common ordered replay, between models trained with different pairings. In the common evaluation, the four-dimensional MEA readings matched element by element in all 4,608 pairs of decision steps. The archived game observation hashes (obs_hash) differed in 288 pairs, and the actions actually selected differed in 301 pairs. Across the 768 pairs of first steps matched by training seed and map, both the archived game observation hashes and the four-dimensional MEA readings matched. In a post hoc diagnostic, the verb probability vectors for each pair of first steps were subtracted element by element, and half the sum of the absolute differences was taken as the total variation distance. The mean was 0.01324, equal to the equally weighted average of the means across training seeds. The actions actually selected, including their targets, matched in 749/768 pairs. The checks relied on the archived registry and per-step evaluation logs. The exact fields serialized for the observation hashes, the tokenized policy prompts, the complete value network inputs, and the intermediate tensors still need to be verified. The archived code set the sampling seed from the map seed and decision index. This implied that the first-step sampling seed settings were identical for the two groups on the common maps. There were no separate per-step records of the actual sampling seeds or random number generator states. The table below lists three comparisons at the six-decision budget. All intervals for the exploration differences crossed zero, leaving the effects of training pairings or input conditions on exploration inconclusive. After six decisions, the two groups with MEA input had higher mean scores and lower mean hydration and satiety reserves than the native R1-only group. In the common evaluation, the interval for the score difference calculated as the intact group minus the shuffled group crossed zero, leaving the effect of training pairings on score inconclusive. In a post hoc analysis, the same set of trajectories from 11/18 was truncated at game tick 6 (*β*=0). The individual 95% intervals for the three exploration differences and the three R1 differences all crossed zero. The hydration differences obtained by subtracting the native R1-only group from each of the two groups with MEA input were below zero, as were their individual 95% intervals. The point estimates for the two satiety differences became positive, and their individual 95% intervals all crossed zero. The reserve results were exploratory.

#### 22554 paired between-group differences at the six-decision budget and reserves at game tick 6

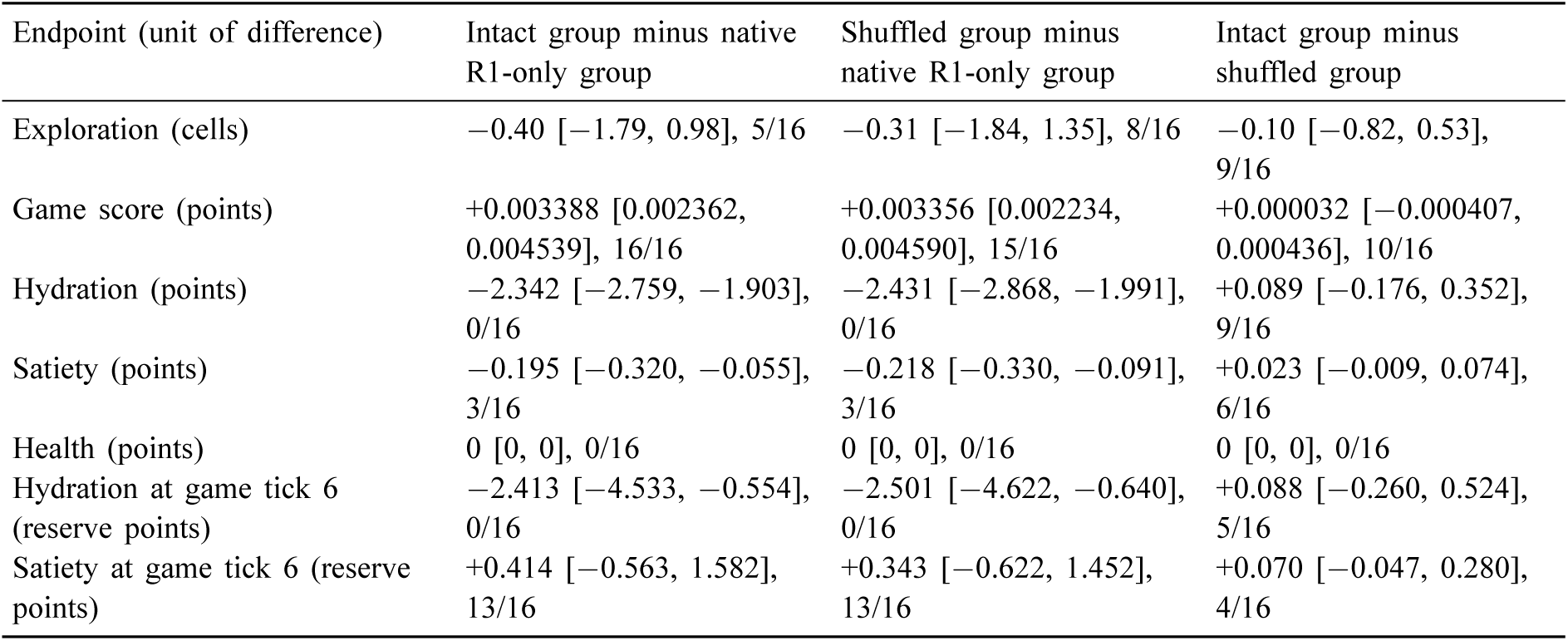

| Endpoint (unit of difference) | Intact group minus native R1-only group | Shuffled group minus native R1-only group | Intact group minus shuffled group |
|---|---|---|---|
| Exploration (cells) | −0.40 [−1.79, 0.98], 5/16 | −0.31 [−1.84, 1.35], 8/16 | −0.10 [−0.82, 0.53], 9/16 |
| Game score (points) | +0.003388 [0.002362, 0.004539], 16/16 | +0.003356 [0.002234, 0.004590], 15/16 | +0.000032 [−0.000407, 0.000436], 10/16 |
| Hydration (points) | −2.342 [−2.759, −1.903], 0/16 | −2.431 [−2.868, −1.991], 0/16 | +0.089 [−0.176, 0.352], 9/16 |
| Satiety (points) | −0.195 [−0.320, −0.055], 3/16 | −0.218 [−0.330, −0.091], 3/16 | +0.023 [−0.009, 0.074], 6/16 |
| Health (points) | 0 [0, 0], 0/16 | 0 [0, 0], 0/16 | 0 [0, 0], 0/16 |
| Hydration at game tick 6 (reserve points) | −2.413 [−4.533, −0.554], 0/16 | −2.501 [−4.622, −0.640], 0/16 | +0.088 [−0.260, 0.524], 5/16 |
| Satiety at game tick 6 (reserve points) | +0.414 [−0.563, 1.582], 13/16 | +0.343 [−0.622, 1.452], 13/16 | +0.070 [−0.047, 0.280], 4/16 |

Note. Both tables draw on the same set of trajectories from the 11/18 replay, with 768 episodes per arm.

For the individual two-sided 95% intervals at the end of six decisions, the set of 48 maps was held fixed and training seeds were resampled 12,000 times. The trajectories were truncated post hoc at game tick 6. The individual two-sided 95% intervals were based on 20,000 iterations of crossed resampling of training seeds and common maps. The count after each interval gives the number of seeds with a within-seed mean difference strictly greater than zero, out of 16 seeds. At game tick 6, the hydration differences for the intact group and the shuffled group, each minus the native R1-only group, were *−*2.413 [*−*4.533, *−*0.554] and *−*2.501 [*−*4.622, *−*0.640] reserve points, respectively. The satiety differences were +0.414 [*−*0.563, 1.582] and +0.343 [*−*0.622, 1.452] reserve points, respectively. The last two rows of the table above give the corresponding counts of seeds with a positive difference. In the direct comparison of the two groups with MEA input, all four individual 95% intervals for hydration and satiety differences across the two budgets crossed zero. All reserve results were exploratory. The basis for a dedicated correction for multiple endpoints still needs to be verified. Health was equal at the end of six decisions. On 11/18, the fixed readout panel failed validation. Before collection of the nine trials began, the registry recorded that the panel had not qualified for real-time training. After collection, exploratory leave-one-block-out classification of the nine trials did not confirm that the stimulation sites could be distinguished. The game replay separately used six trials for training and retained three trials, as described in Section S4.2. A separate evaluation on 11/19 used holdouts by stimulation-site file and two budgets of 48 and 192 game ticks. The exploration comparison intervals crossed zero at both of the two budgets. Appendix B gives the individual results, training rewards, and reused checkpoints.

Exploratory paired results for the same batch of traces on chip 22554 at game tick 6. We selected the six-decision traces from 11/18 post hoc, with *β*=0. Each arm included 16 seeds *×* 48 maps, for a total of 768 games. We reported paired mean differences, two-sided 95% intervals for individual comparisons, and counts of favorable seeds. We obtained the intervals by crossed resampling of seeds and common maps over 20,000 resamples. A seed was favorable if its mean difference across maps was strictly greater than zero. The analysis of reserve endpoints at that game tick was exploratory. The archived nominal 98.333% intervals at the end of the table use the same interval level as the original reward-weight sweep. The basis for a multiplicity correction specific to the family of reserve endpoints at game tick 6 remains to be verified.

| Endpoint and interval level | Intact group minus native R1-only baseline | Shuffled group minus native R1-only baseline | Intact group minus shuffled group |
|---|---|---|---|
| Exploration (cells per game), 95% per comparison | −1.98 [−5.56, 1.38], 2/16 | −1.95 [−5.52, 1.43], 3/16 | −0.03 [−0.68, 0.58], 8/16 |
| R1 (points per game), 95% per comparison (the last two rows of the preceding table report paired differences, intervals, and counts of seeds with positive differences for hydration and satiety at the same game tick) | +0.000842 [−0.000633, 0.003897], 8/16 | +0.000832 [−0.000642, 0.003832], 8/16 | +0.000010 [−0.000569, 0.000594], 11/16 |
| Exploration (cells per game), archived 98.333% | [−6.38, 2.13] | [−6.32, 2.12] | [−0.85, 0.75] |
| R1 (points per game), archived 98.333% | [−0.000748, 0.004850] | [−0.000753, 0.004899] | [−0.000862, 0.000882] |
| Hydration (reserve points), archived 98.333% | [−5.088, −0.201] | [−5.170, −0.318] | [−0.376, 0.670] |
| Satiety (reserve points), archived 98.333% | [−0.769, 1.888] | [−0.860, 1.758] | [−0.094, 0.352] |

### S5.6 Low-health, severe-thirst, and severe-hunger starts for the second specimen on chip 22021

The low-health start began with 5 health points and 95 points each for hydration and satiety. By game tick 60, 2,048/2,048 games in each arm had survived, and survival reached its ceiling. The draft specified the number of newly revealed cells at the cutoff tick or at earlier death as the primary endpoint. The intact group, all-zero group, and shuffled group averaged 131.72, 126.125, and 129.44 cells, respectively. The intact group minus the all-zero group yielded a difference of +5.59 cells (99.167% interval [*−*0.19, 11.26], with 25/32 favorable seeds). The intact group minus the shuffled group yielded a difference of +2.28 cells (interval [*−*1.92, 7.13], with 21/32 favorable seeds). The two intervals crossed zero, leaving the effect on exploration under the low-health start inconclusive.

The other two starts began with 25 health points. The severe-thirst start began with 1.6 hydration points, and the severe-hunger start began with 4 satiety points. Each arm included 2,048 games per scenario. Under severe thirst, the intact group, all-zero group, and shuffled group survived in 803, 862, and 837 games, respectively. Under severe hunger, the corresponding counts were 1,210, 1,275, and 1,233. In the two scenarios, the point estimates of the survival-rate differences for the intact group minus each of the two control groups were all negative (Table 2). All four full-sample 99.167% intervals crossed zero. For the intact group minus the all-zero group and minus the shuffled group, in that order, the upper bounds were +2.73 and +2.83 percentage points under severe thirst and +2.34 and +3.29 percentage points under severe hunger. A post hoc analysis retained only the 24 newly added seeds and reused the original 64 pilot maps. All four intervals in that analysis also crossed zero, with upper bounds of +3.45, +2.86, +2.86, and +3.19 percentage points in the same order. Every upper bound in both analyses was below the value of +5 percentage points used in the power simulation. Smaller improvements and decreases in survival rate remained compatible with these intervals. A review against the available registration draft on all 64 reused pilot maps, including the analysis of exploration under the low-health start, found that none of the six primary comparisons met the criteria (0/6) and none of the three scenarios met the criteria (0/3). Whether a single protocol was locked before unblinding and whether the complete set of receipts was sealed and archived both remain to be independently verified. Independent validation on new maps has yet to be conducted.

Restricted waiting time used all games as the denominator. For each legal drinking or eating action, the trajectory was checked tick by tick. Success was recorded at the first game tick at which hydration or satiety exceeded its value at the start of the action. Actions that extended beyond game tick 60 were checked only through game tick 60, inclusive. Waiting time was the number of game ticks from the start of the game to the first success. Games in which the player died before the first success or in which no success had occurred by game tick 60 were assigned a waiting time of 60 game ticks. By definition, a success at game tick 60 also counted as 60 game ticks. A waiting time of 60 alone therefore cannot determine whether the game had a success. Differences were calculated by subtracting the intact group from the control group, with positive values indicating a shorter mean restricted waiting time in the intact group. For rehydration under the severethirst start, the differences relative to the all-zero group and the shuffled group were *−*1.54 [*−*4.26, 0.91] and *−*0.74 [*−*2.78, 1.15] game ticks, respectively, with 7/32 and 13/32 seeds favoring the intact group. For eating under the severe-hunger start, the corresponding differences were *−*0.78 [*−*2.73, 1.18] and *−*0.32 [*−*2.06, 1.25] game ticks, with 10/32 and 13/32 seeds favoring the intact group, respectively. The full-sample point estimates for all four restricted waiting time differences were negative, and all two-sided 98.75% adjusted intervals crossed zero. On rechecking under the archived procedure described in Section S3.6, 0/4 comparisons met the criteria. In a post hoc analysis that retained only the 24 additional seeds and reused all 64 pilot maps, all four intervals also crossed zero (Section S8). The effects on waiting for rehydration or eating remained inconclusive. According to the archived game-by-game verification summary, the rehydration success flags in these severe-thirst games and the eating success flags in these severe-hunger games each agreed with the survival flags at game tick 60. There were 2,048 games per group per scenario, and the discrepancy count was 0 in each of the six scenario-by-group combinations. According to the summary of the cross-check between archived action events and historical native replays, there were 12,288 games across the three groups and two start conditions, with rehydration assessed under severe thirst and eating under severe hunger. Of these, 6,220 games had their first success before game tick 60, and 0 games had their first success exactly at game tick 60. In the remaining 6,068 games, the player died before the cutoff without a corresponding success event. Each of those games was assigned a restricted waiting time of 60 game ticks. Among games with an earlier first success, 127 had another success at game tick 60. Their restricted waiting time was based on the earlier first success. Of the 2,048 games per arm under the severe-hunger start, the numbers of games with a first successful eating event were 1,210, 1,275, and 1,233 for the intact, all-zero, and shuffled groups, respectively (59.08%, 62.26%, and 60.21%). Among successful games, the mean game ticks of the first successful eating event were 16.32, 17.29, and 16.61, respectively. These three conditional means can be recovered by multiplying each group’s unrounded mean restricted waiting time across all games by 2,048, subtracting the number of unsuccessful games multiplied by 60, and dividing by the number of successful games. The intact group had a longer mean restricted waiting time across all games and a lower proportion of successful games. Among successful games, eating occurred earlier on average in the intact group. Successful games in each arm were selected from that arm’s own trajectories. Games were matched post hoc by training seed and map, yielding 2,048 pairs per comparison. In the order of success in both games, success only in the intact group, success only in the control group, and success in neither game, the counts were 1,081, 129, 194, and 644 pairs for the intact group versus the all-zero group and 1,104, 106, 129, and 709 pairs for the intact group versus the shuffled group. All differences below were computed by subtracting the intact group from the all-zero group and from the shuffled group separately. Among the 1,081 and 1,104 pairs with success in both games, the mean differences in time to the first successful eating event were +0.068 and +0.152 game ticks per pair, respectively, with corresponding net differences of +74 and +168 game ticks. In pairs with no success in either game, both waiting times were 60 game ticks, yielding a net difference of zero. Pairs with success on only one side contributed *−*1,676 and *−*815 game ticks, respectively. Across all 2,048 pairs, the resulting net differences in restricted waiting time were *−*1,602 and *−*647 game ticks, respectively. Restricted waiting time across all games was a secondary endpoint in the draft registration, and both of the two adjusted 98.75% intervals crossed zero. The time differences among pairs with success in both games described only games selected by outcome. The effect on whether eating occurred sooner remained inconclusive. Eating success and survival agreed game by game in this scenario, so the two outcomes cannot independently corroborate that effect. These measures described observable survival and replenishment behavior. Subjective fear was not measured.

## S6 Analysis

### S6.1 Reanalysis at a Fixed Game Tick

Any one navigation action could span multiple game ticks. In a post hoc exploratory analysis of the same six-decision trajectories from Section S5.1, we replayed the recorded actions and truncated the replay at game tick 6. If an action extended past the cutoff, we read the map from the trajectory state at game tick 6. The all-zero group, intact group, and unstimulated group averaged 34.26, 35.48, and 35.80 newly revealed cells per episode, respectively. In paired comparisons with the all-zero group, the intact group revealed 1.21 more cells and the unstimulated group revealed 1.53 more cells. The proportions of seeds with a difference in the specified direction, positive in both comparisons, were 13/16 and 16/16, respectively. The 95% intervals from crossed resampling of seeds and maps [11] were [0.08, 2.59] and [0.14, 3.18], respectively. A post hoc sensitivity analysis used the nominal Bonferroni level of 98.333% for the three nonzero *β* comparisons in the original *β* sweep. The two archived crossed-resampling intervals were [*−*0.16, 2.97] and [*−*0.12, 3.65] cells per episode, respectively. Both crossed zero. When the estimand allowed the maps to vary, the exploration difference at game tick 6 remained inconclusive. In the same trajectories, the exploration difference at game tick 6, calculated as the intact group minus the unstimulated group, was *−*0.32 cells per episode. The 95% interval was [*−*1.20, 0.48], the 98.333% interval was [*−*1.43, 0.67], and the difference was positive for 6/16 seeds. The timing decomposition in Section S5.2 was limited to the first-day subtrial. After pooling the initial and expansion batches, on the 48 shared maps, the mean differences in R1 at the end of six decisions were +0.004065 and +0.004339 points per episode for the intact group minus the all-zero group and the unstimulated group minus the all-zero group, respectively. In each of the two comparisons, 16/16 training seeds had a positive mean difference. With these maps fixed and only training seeds resampled, the per-comparison 95% intervals were [0.002794, 0.005493] and [0.003268, 0.005497] points per episode, respectively. At game tick 6, the mean differences in R1 for the intact group minus the all-zero group and the unstimulated group minus the all-zero group were +0.000443 and +0.000220 points per episode, respectively. The per-comparison 95% crossed-resampling intervals were [*−*0.000022, 0.001412] and [*−*0.000144, 0.000884] points per episode, respectively. Both crossed zero. The latter set of intervals reflected variability from changing maps under the same map-generation mechanism and from changing training seeds.

#### Exploration-difference intervals for the first specimen on chip 22021, compared by estimand (cells per episode)

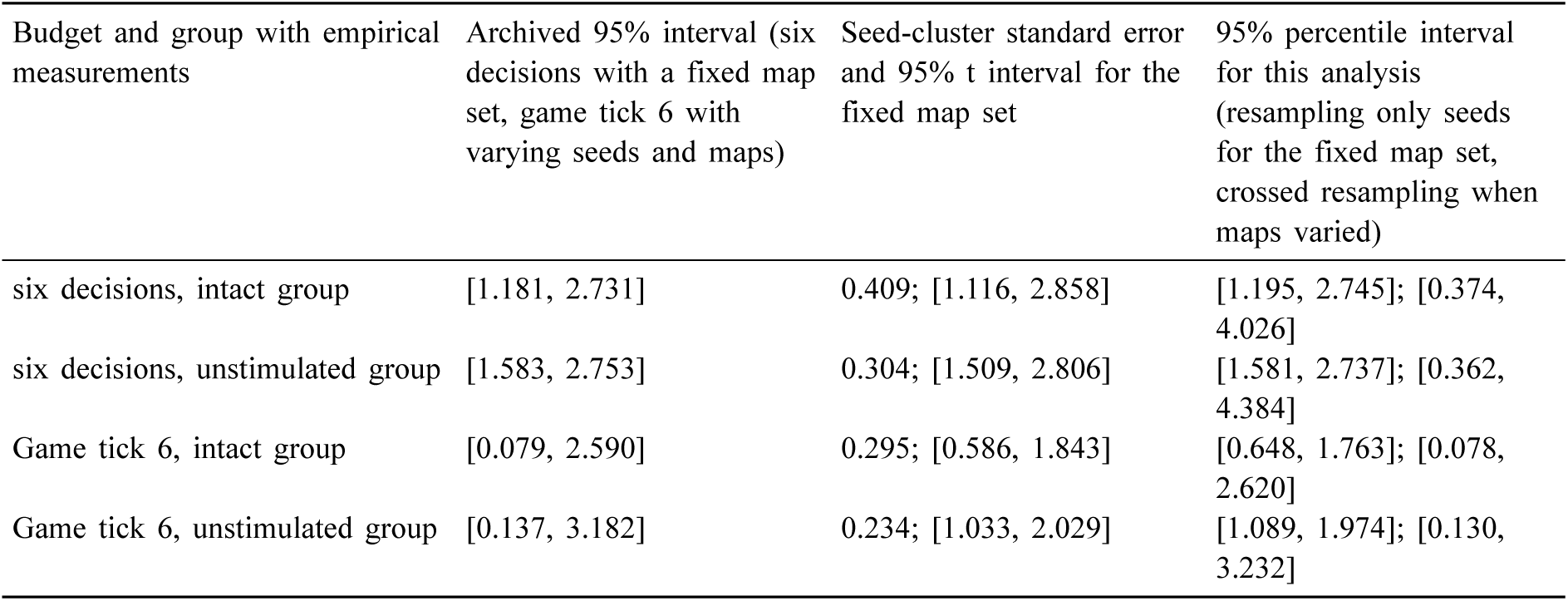

Each difference was calculated by subtracting the all-zero group’s result from the result of the group using measured neural input. The supplementary interval analysis used 40,000 paired resamples and a random seed of 2026112201. Clustered standard errors were calculated using 16 training seeds, and the t intervals used 15 degrees of freedom. The post hoc sensitivity analysis at game tick 6 with *β*=0 retained the nominal Bonferroni level of 98.333% for the original three nonzero *β* comparisons. With the original 48 maps fixed and only seeds resampled, the percentile intervals were [0.522, 1.876] cells per episode for the intact group minus the all-zero group and [1.000, 2.068] cells per episode for the unstimulated group minus the all-zero group. The corresponding percentile intervals from crossed resampling of seeds and shared maps were [*−*0.152, 3.017] and [*−*0.158, 3.692] cells per episode. At the same nominal level, the t intervals with maps fixed and clustering by seed were [0.420, 2.009] and [0.902, 2.161] cells per episode, respectively. The lower bounds of all intervals from the two fixed-map methods were above zero. The two intervals from crossed resampling of seeds and maps both crossed zero. The exploration difference at equal elapsed game time remained inconclusive for the estimand that allowed maps to vary.

### S6.2 Per-Episode Distribution

At the end of six decisions, among 768 combinations matched by training seed and map, the intact group tied with the all-zero group in 615 pairs, and the unstimulated group tied with the all-zero group in 602 pairs. All three groups had the same number of newly revealed cells in 577 pairs. Of the 48 maps tested, 14 met the following condition at every one of the 16 training seeds. For each measured group, subtracting the allzero group’s count of newly revealed cells from that group’s count gave a difference of zero. Equal counts of newly revealed cells could still correspond to different locations or routes. For each comparison, the 768 differences were sorted in ascending order, and 5% was removed from each end (rounded down to 38 pairs per end). The mean differences across the remaining 692 pairs were +1.079 and +1.172 cells, respectively. Removing only the 38 largest positive differences and retaining all negative differences left 730 pairs. The post-trimming mean differences reported in Section S5.1 describe only these 730 pairs and indicate that a small number of larger positive differences influenced the original means.

Both of the two trimming procedures ranked pairs by their differences and removed a fixed number of pairs. When differences were tied at the cutoff, the choice of which tied pair to remove did not change the posttrimming mean.

At game tick 6, in the same set of 768 trajectory pairs, the win/tie/loss counts were 79/654/35 for the intact group minus the all-zero group and 88/644/36 for the unstimulated group minus the all-zero group. Counts of newly revealed cells were identical across all three groups in 622 pairs. Of the 48 maps tested, the count was identical across all three groups at every one of the 16 seeds on 14 maps at the end of six decisions and on 20 maps at game tick 6. At game tick 6, the win rates across all pairs in the two comparisons were 10.29% and 11.46%, respectively. Both median differences were zero, and the mean differences expressed relative to the all-zero group’s 34.264 cells were +3.55% and +4.47%, respectively. The paired dz values were 0.166 and 0.197, and the paired rank-biserial correlations for the nonzero differences were 0.463 and 0.546, respectively. After the differences were sorted and 38 pairs were removed from each end, the mean differences across the remaining 692 pairs were +0.539 and +0.705 cells. When only the 38 largest positive differences were removed, the mean differences across the remaining 730 pairs were *−*0.138 and +0.095 cells. These means describe the games retained after trimming by outcome direction. In a post hoc influence analysis by whole map, M29 contributed 414/1,526 cells (27.1%) and 520/1,657 cells (31.4%), respectively, to the signed net differences in the two comparisons at the end of six decisions. At game tick 6, its contributions were 238/933 cells (25.5%) and 320/1,176 cells (27.2%), respectively. After M29 was excluded, each comparison at game tick 6 retained 752 pairs across the remaining 47 maps, with mean differences of +0.924 and +1.138 cells per game, respectively. At game tick 6, the net difference for the unstimulated group minus the all-zero group was largest on M45, at 368 cells. After that map was excluded, the mean difference was +1.074 cells per game. Together, M29 and M45 accounted for 49.6% and 58.5% of the observed net differences in the two comparisons at game tick 6, respectively. When both maps were excluded post hoc based on the net differences observed in this set, all 16 seeds were retained on each of the remaining 46 maps, yielding 736 pairs per comparison. The descriptive mean differences were +0.639 and +0.663 cells per game for the intact group minus the all-zero group and the unstimulated group minus the all-zero group, respectively. At game tick 6, both the comparison-specific 95% intervals and the nominal 98.333% intervals were calculated using data from the original 48 maps. Appendix A reports the mean differences by map and the ranges of mean differences obtained by excluding each map in turn.

### S6.3 Power for the next round

We used a pilot paired matrix of 8 seeds *×* 64 maps to simulate a future design with 32 seeds *×* 64 maps. For each of the two sets of paired differences at each endpoint, we subtracted the corresponding pilot mean and added the specified true improvement. For each of the 4,000 outer samples, we used crossed resampling to draw 40,000 resamples. All comparisons shared the same seed and map indices. Table 4 retains the earlier simulation and the paired simulation. In the paired simulation, the detection probabilities for true increases of 5 and 8 in the count of newly revealed cells from a low-health start were 12.80% and 71.73%, respectively. With a true improvement of 5 percentage points in survival, the detection probabilities for the severe-thirst and severe-hunger starts were 14.15% and 6.30%, respectively. Success required meeting both the adjusted interval criterion and the threshold of at least 24/32 seeds with a difference in the specified direction against each of the two controls at the same endpoint. We did not estimate the probability of all six comparisons across the three endpoints meeting the criteria.

**Table 4.**
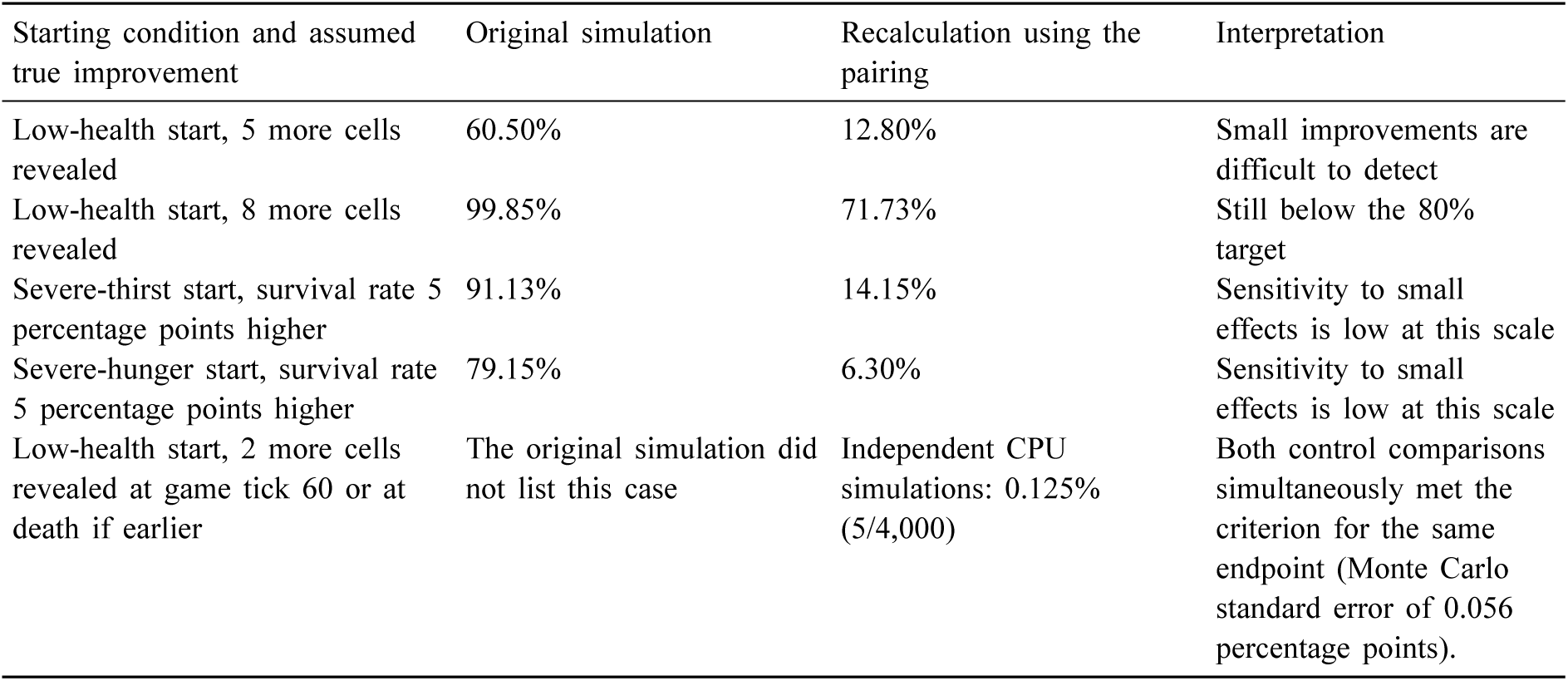
Simulated probability of simultaneously outperforming the two controls with 32 seeds *×* 64 maps. In the original paired simulation, when the true improvement in low-health exploration was 5 cells, the two control comparisons jointly met their criteria in 512/4,000 runs (12.80%). In the separately calculated +2-cell scenario, the corresponding result was 5/4,000 (0.125%, with a Monte Carlo standard error of 0.056 percentage points). Each probability applies to joint success against the two controls at the same endpoint.

| Starting condition and assumed true improvement | Original simulation | Recalculation using the pairing | Interpretation |
|---|---|---|---|
| Low-health start, 5 more cells revealed | 60.50% | 12.80% | Small improvements are difficult to detect |
| Low-health start, 8 more cells revealed | 99.85% | 71.73% | Still below the 80% target |
| Severe-thirst start, survival rate 5 percentage points higher | 91.13% | 14.15% | Sensitivity to small effects is low at this scale |
| Severe-hunger start, survival rate 5 percentage points higher | 79.15% | 6.30% | Sensitivity to small effects is low at this scale |
| Low-health start, 2 more cells revealed at game tick 60 or at death if earlier | The original simulation did not list this case | Independent CPU simulations: 0.125% (5/4,000) | Both control comparisons simultaneously met the criterion for the same endpoint (Monte Carlo standard error of 0.056 percentage points). |

We recomputed the simulations using an independent CPU random stream. True improvements of 5 and 8 cells from a low-health start were detected in 494/4,000 (12.350%) and 2,856/4,000 (71.400%) samples, respectively. With a true improvement of 5 percentage points, the severe-thirst and severe-hunger starts yielded detections in 533/4,000 (13.325%) and 256/4,000 (6.400%) samples, respectively. In the additional +2-cell scenario for exploration from a low-health start, the intact group met the criteria against the all-zero group 84 times and against the shuffled group 92 times. The two comparisons both met the criteria in 5/4,000 samples (0.125%). The Monte Carlo standard error of the joint detection rate was 0.056 percentage points, and the Wilson 95% Monte Carlo interval ranged from 0.0534% to 0.2923%. The estimates were conditional on the pilot matrix and the location-shift assumption. The survival-rate simulation shifted the paired differences in proportions directly. The assumed improvement of +2 cells by game tick 60 from a low-health start differed in budget and controls from the observed mean difference over six decisions from a normal start.

The additive normal simulation summed two mutually independent standard normal effects, one for seed and one for map, each with mean zero and variance 1. The interaction was set to zero. For each setting, 2,048 datasets were simulated in 64 independent batches. The 32 datasets in each batch shared the resampling weights used to construct the percentile intervals. Each percentile interval used 40,000 resamples. The target for the fixed map set was the mean effect of the maps drawn in that replicate. The population target when both seeds and maps were changed was zero. The Monte Carlo standard error of coverage was calculated as the sample standard deviation of the batch coverage proportions divided by 8. In the additive normal simulation with 16 seeds *×* 48 maps and nominal 95% coverage, seed-only percentile intervals had higher coverage for the fixed-map mean than for the population mean when both seeds and maps were changed. For the population mean, percentile intervals from crossed resampling had higher coverage than seed-only percentile intervals. A separate simulation used the 16 seeds *×* 48 maps matrix of paired differences in the number of newly revealed cells from the first specimen on chip 22021 as a discrete empirical population. It retained zero differences, ties, and seed-by-map interactions. Each scenario had 4,096 independent outer samples, and each percentile interval used 40,000 resamples. With the original map set fixed, only seeds were resampled at the outer level, and the target was the mean for the original map set. When both seeds and maps were changed, seed rows and map columns were resampled separately at the outer level, and the target was the mean of the full empirical matrix. In this simulation, each of these two targets equaled the equally weighted mean of the corresponding original matrix. For the intact group minus the all-zero group and the unstimulated group minus the all-zero group, respectively, the empirical true values at the end of six decisions were +1.987 and +2.158 cells per game. At game tick 6, they were +1.215 and +1.531 cells per game. For samples with resampled maps, a conditional true value could also be calculated using the map positions drawn in that replicate and all original seeds, retaining duplicate map positions. The Monte Carlo standard errors for the empirical matrix were calculated from the coverage indicators of the independent outer samples. In both coverage tables, *±* denotes one Monte Carlo standard error, in percentage points. The table “Coverage for the Empirical Matrix of Paired Differences from the First Specimen” reports the hit counts, coverage rates, and standard errors for the nominal 95% simulation at the end of six decisions. In each of the two comparisons at the end of six decisions, seed-only percentile intervals had lower coverage of the full-matrix mean in samples with resampled maps than the corresponding coverage with the original map set fixed. Their coverage of the full-matrix mean was also lower than the coverage of the respective conditional true values by the same intervals in the same samples with resampled maps. With the full-matrix mean as the target, percentile intervals from crossed resampling had higher coverage than both seed-only percentile intervals and seed-cluster t intervals in each of the two comparisons. The same trajectories were retrospectively truncated at game tick 6. At the nominal 98.333% level, in the order intact group minus all-zero group and unstimulated group minus all-zero group, coverage for seed-only percentile intervals on the fixed map set was 96.19% and 96.95%, respectively, with 3,940 and 3,971 hits and standard errors of 0.30 and 0.27 percentage points. Coverage for percentile intervals from crossed resampling of both seeds and maps was 98.02% and 96.44%, respectively, with 4,015 and 3,950 hits and standard errors of 0.22 and 0.29 percentage points. Coverage in the two simulations applied only to the respective generating models and estimation targets. Interval coverage under the actual map mechanism and on independent biological recordings remains unknown.

#### Coverage in the General Additive Normal Simulation

| Sample size and estimand | Nominal level and interval method | Number of intervals covering the target / number of simulated datasets | Coverage ± Monte Carlo standard error (percentage points) |
|---|---|---|---|
| 16 seeds × 48 maps, mean for the fixed map set | 95%, seed percentile interval | 1,908/2,048 | 93.16% ± 0.56 |
| 16 seeds × 48 maps, mean for the fixed map set | 95%, seed-cluster t interval | 1,966/2,048 | 96.00% ± 0.43 |
| 16 seeds × 48 maps, population mean over varying seeds and maps | 95%, seed percentile interval | 1,803/2,048 | 88.04% ± 0.69 |
| 16 seeds × 48 maps, population mean over varying seeds and maps | 95%, percentile interval from crossed resampling | 1,921/2,048 | 93.80% ± 0.60 |
| 32 seeds × 64 maps, population mean over varying seeds and maps | 99.167%, percentile interval from crossed resampling | 2,028/2,048 | 99.02% ± 0.20 |

#### Coverage for the Empirical Matrix of Paired Differences from the First Specimen

| Budget and estimand | Nominal level and method | Intact group minus all-zero group | Unstimulated group minus all-zero group |
|---|---|---|---|
| Empirical matrix after six decisions, empirical mean with the original map set held fixed | 95%, seed percentile method | 3,805/4,096, 92.90%±0.40 | 3,812/4,096, 93.07%±0.40 |
| Empirical matrix after six decisions, empirical mean with the original map set held fixed | 95%, seed-cluster t method | 3,884/4,096, 94.82%±0.35 | 3,904/4,096, 95.31%±0.33 |
| Empirical matrix after six decisions, full-matrix mean with new seeds and maps | 95%, seed percentile method | 2,681/4,096, 65.45%±0.74 | 2,154/4,096, 52.59%±0.78 |
| Empirical matrix after six decisions, full-matrix mean with new seeds and maps | 95%, crossed percentile method | 3,860/4,096, 94.24%±0.36 | 3,812/4,096, 93.07%±0.40 |
| Empirical matrix after six decisions, full-matrix mean with new seeds and maps | 95%, seed-cluster t method | 2,877/4,096, 70.24%±0.71 | 2,382/4,096, 58.15%±0.77 |
| Empirical matrix at game tick 6, empirical mean with the original map set held fixed | 95%, seed percentile method | 3,784/4,096, 92.38%±0.41 | 3,789/4,096, 92.50%±0.41 |
| Empirical matrix at game tick 6, empirical mean with the original map set held fixed | 95%, seed-cluster t method | 3,881/4,096, 94.75%±0.35 | 3,879/4,096, 94.70%±0.35 |
| Empirical matrix at game tick 6, full-matrix mean with new seeds and maps | 95%, seed percentile method | 2,818/4,096, 68.80%±0.72 | 2,165/4,096, 52.86%±0.78 |
| Empirical matrix at game tick 6, full-matrix mean with new seeds and maps | 95%, seed-cluster t method | 3,022/4,096, 73.78%±0.69 | 2,391/4,096, 58.37%±0.77 |
| Empirical matrix at game tick 6, full-matrix mean with seeds and maps varied | 95%, crossed percentile | 3,884/4,096, 94.82%±0.35 | 3,807/4,096, 92.94%±0.40 |
| Empirical matrix at game tick 6, empirical mean with the original map set held fixed | 98.333%, seed-cluster t | 4,016/4,096, 98.05%±0.22 | 4,021/4,096, 98.17%±0.21 |
| Empirical matrix at game tick 6, full-matrix mean with seeds and maps varied | 98.333%, seed percentile | 3,148/4,096, 76.86%±0.66 | 2,551/4,096, 62.28%±0.76 |
| Empirical matrix at game tick 6, full-matrix mean with seeds and maps varied | 98.333%, seed-cluster t | 3,398/4,096, 82.96%±0.59 | 2,845/4,096, 69.46%±0.72 |

## S7 Discussion

The six-decision comparison for the first specimen on chip 22021 had no shuffled group. The other two experiments used different shuffling procedures. On chip 22554, the intact group and the shuffled group used different trial pairings during training. Both preserved the order of the preand post-stimulation windows within any one trial. The two groups shared ordered replay during evaluation. The four-dimensional MEA inputs were identical component by component for paired decisions. For some decisions, the archived game observation summary (obs_hash) or the action actually selected differed. According to the archived-log verification summary cited in Section S5.5, all 768 first-step pairs under shared ordered replay matched on the initial game observation summary and the four-dimensional MEA readings. Under the first-step pairing criteria in Section S5.5, the mean total variation distance between the verb probability distributions was 0.01324. The actions actually selected, including their targets, matched in 749/768 pairs. The first-step matches were verified against the archived registry and step-by-step evaluation logs. Under the archived code’s rules for setting sampling seeds from the map seed and decision index, the two groups had identical first-step sampling seed settings on shared maps. The sampling seeds actually used during the runs and the random number generator states at the time lacked separate records. Later observations could diverge depending on earlier actions. The selected actions and targets show only what was actually chosen. They cannot be used to reconstruct the probabilities of all targets under each verb. This post hoc comparison cannot identify an effect specific to organoid signals or separate the effects of supplying measured readings during training and evaluation for the first specimen on chip 22021. Under this shared replay condition, the exploration difference over six decisions between the two training pairings (intact group minus shuffled group) was *−*0.10 cells (95% interval [*−*0.82, 0.53], positive for 9/16 seeds). For the low-health start of the second specimen on chip 22021, the difference at game tick 60 or at death was +2.28 cells (99.167% interval [*−*1.92, 7.13], positive for 21/32 seeds). The two intervals crossed zero. Whether the difference in training pairings on chip 22554 corresponded to an exploration effect remained inconclusive, as did whether the difference in block order on chip 22021 corresponded to an exploration effect. The shuffled group for the second specimen on chip 22021 preserved the order within two-second blocks. According to the summary of step-by-step checks against archived records, all training seeds in the shuffled group shared one window order. During evaluation, 64 complete permutations were assigned by map, and four distinct four-dimensional vectors of measured readings were recorded at the first step across maps. For a given map and game tick, the assigned window was reused across training seeds and starting scenarios (see Section S3.4). The paired difference for the low-health start describes the overall performance difference under the two training and evaluation replay arrangements. The effects of continuity between blocks, the correspondence between individual decisions and recorded segments, changes in replay from training to evaluation, and the assignment of evaluation permutations by map could not be estimated separately. Whether the policy used the input differences assigned by map remains unknown.

On chip 22021, each group used its own input-reading procedure for both training and evaluation in the sixdecision experiment with the first specimen and in the stress trial with the second specimen. Paired differences for the same seed and map cannot distinguish action preferences acquired during training from the frozen model’s response to the current step’s readings at inference. Setting the model’s neural input to zero in the all-zero group simultaneously removed differences in magnitude, variability, correlation structure, and temporal variation in the four readings. Generic time-varying inputs could still explain the observed group differences. The existing comparisons cannot isolate a contribution specific to the organoid source. The firststep diagnostic in Section S5.5 was limited to the outputs, under shared ordered replay, of the policies obtained from the two training pairings on chip 22554. Future work could cross the assignment of training and evaluation input-reading procedures in the chip 22021 design, independently randomize the assignment of multiple complete block permutations within each map, and compare action probabilities at matched game states. Holding the evaluation input-reading procedure fixed would allow comparisons between training pairings. Holding the training input-reading procedure fixed would allow comparisons between different evaluation permutations within the same map, with differences between permutations also reported for each map. With the model and initial game state fixed on the same map, the replay start could be shifted within a continuous recording while preserving the original block adjacency to compare the alignment of segments with game ticks. In repeated games on the same map, the marginal distributions of assigned segments and withinblock positions could be matched at each prespecified game tick. The condition that preserves the original block order could then be compared with the condition that rearranges adjacent blocks. For both the replaystart shift and the block-order comparison, full-game outcomes would be summarized by assignment scheme. The game ticks actually visited and the windows actually read would be saved at each step to check whether action durations caused segment alignment and inter-block adjacency to continue varying together in the inputs actually read. If the two manipulations still overlap in the inputs actually read, estimation would be limited to the combined difference between the corresponding schemes. A separate four-dimensional synthetic input condition could match the per-dimension amplitude distributions, rates of change between adjacent windows, and cross-dimensional correlations of the measured recordings. This condition could be compared separately with count-based exploration rewards, random network distillation (RND) [15], tuning entropy regularization alone, and NoisyNet. These controls have yet to be run. Follow-up validation of the severe-hunger start could use independent recordings and new maps, prespecify the cumulative incidence of first successful eating, treat death before first eating as a competing event, and continue to report restricted waiting time across all games. Paired timing differences would remain descriptive for pairs in which both games included successful eating. The plan for future independent validation is to use maps that were unused in the pilot, independently seal the endpoints, thresholds, run design fingerprint, and receipt list before viewing the evaluation results, and report whether the criteria were met individually and overall under the same draft criteria reviewed in this study.

Electrode readings can indirectly affect R1 through the selected action and can also change the advantage used for policy updates through value estimation. The two input pathways were handled simultaneously in each group, and their individual effects still could not be separated.

## S8 Limitations

### Written registration and actual execution of the adverse-start trials

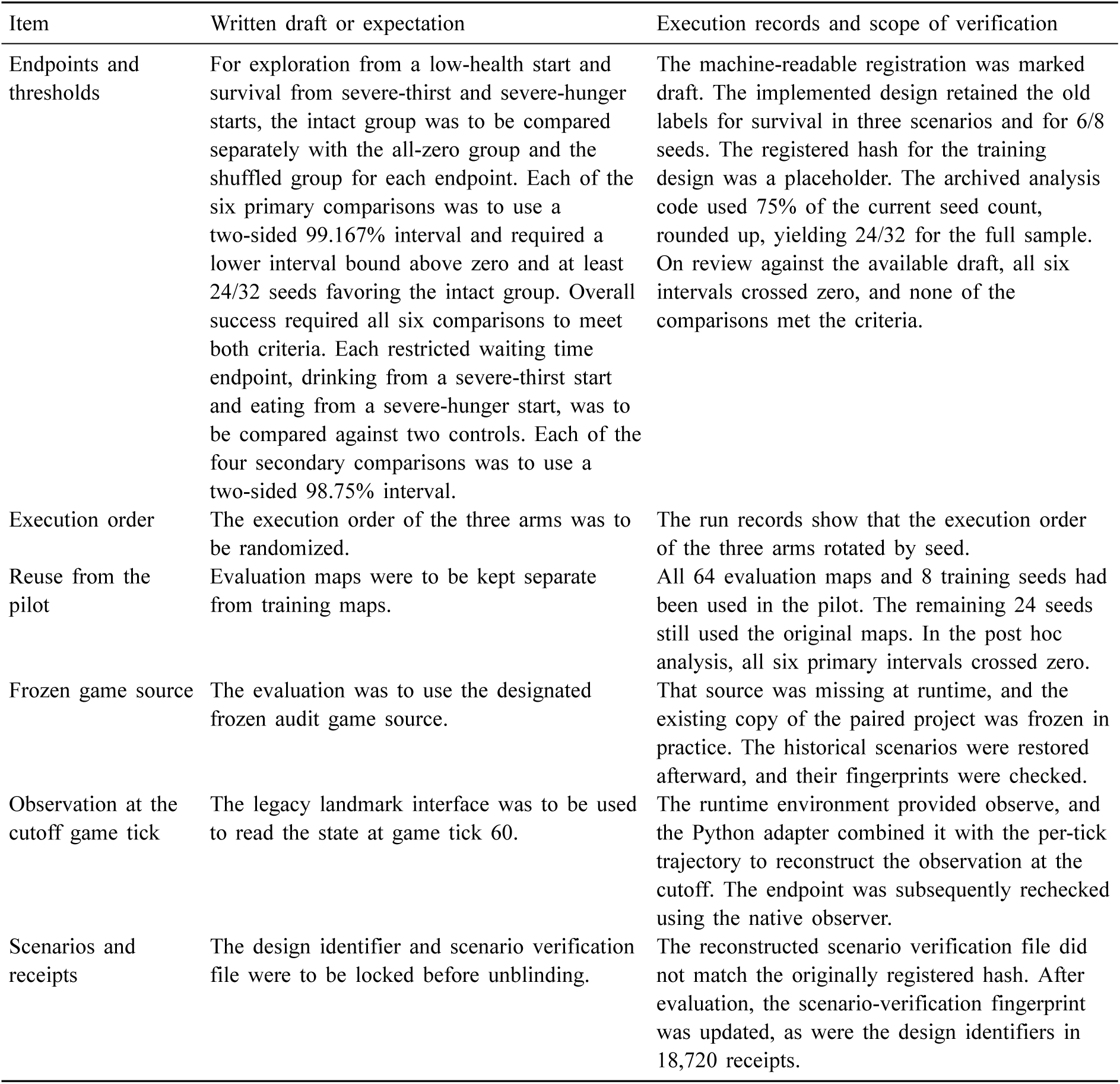
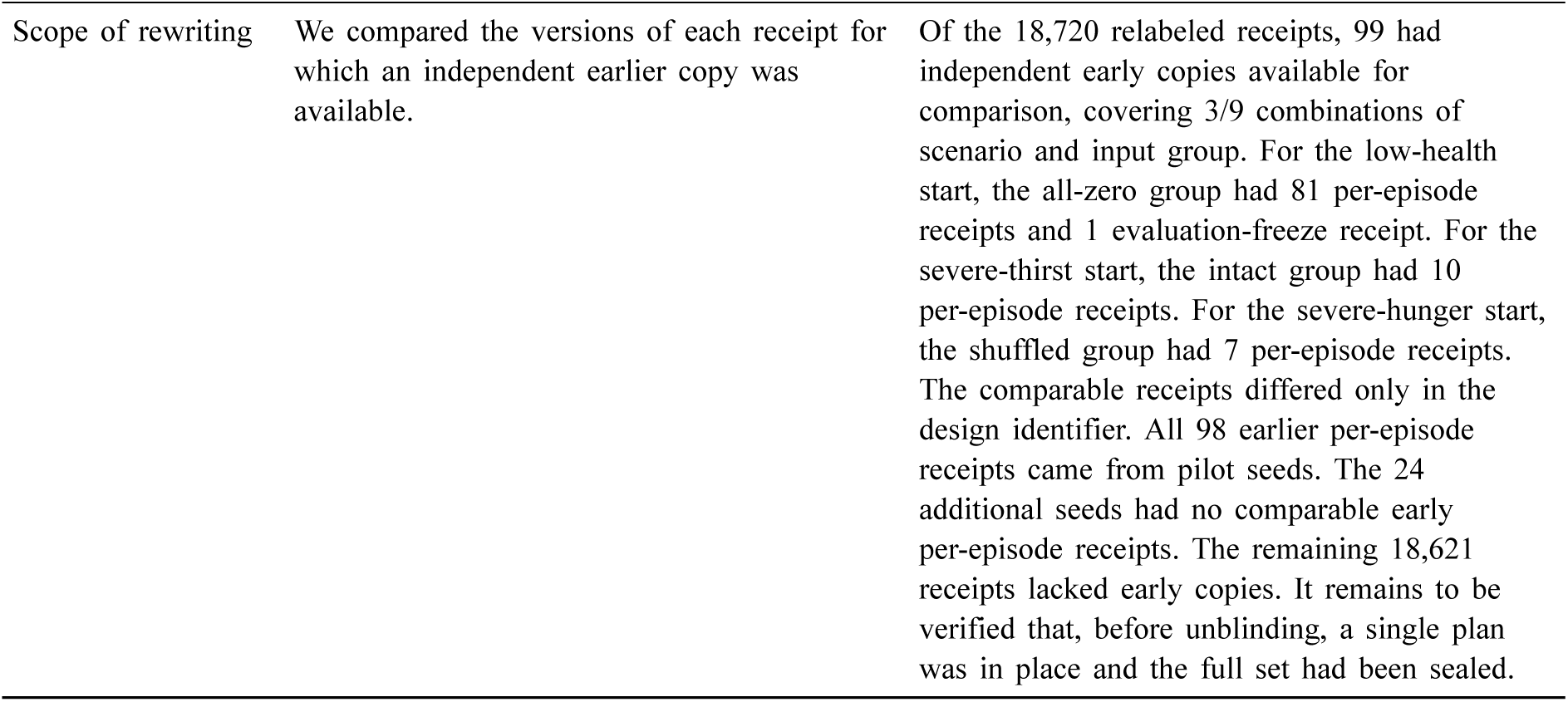

| Item | Written draft or expectation | Execution records and scope of verification |
|---|---|---|
| Endpoints and thresholds | For exploration from a low-health start and survival from severe-thirst and severe-hunger starts, the intact group was to be compared separately with the all-zero group and the shuffled group for each endpoint. Each of the six primary comparisons was to use a two-sided 99.167% interval and required a lower interval bound above zero and at least 24/32 seeds favoring the intact group. Overall success required all six comparisons to meet both criteria. Each restricted waiting time endpoint, drinking from a severe-thirst start and eating from a severe-hunger start, was to be compared against two controls. Each of the four secondary comparisons was to use a two-sided 98.75% interval. | The machine-readable registration was marked draft. The implemented design retained the old labels for survival in three scenarios and for 6/8 seeds. The registered hash for the training design was a placeholder. The archived analysis code used 75% of the current seed count, rounded up, yielding 24/32 for the full sample. On review against the available draft, all six intervals crossed zero, and none of the comparisons met the criteria. |
| Execution order | The execution order of the three arms was to be randomized. | The run records show that the execution order of the three arms rotated by seed. |
| Reuse from the pilot | Evaluation maps were to be kept separate from training maps. | All 64 evaluation maps and 8 training seeds had been used in the pilot. The remaining 24 seeds still used the original maps. In the post hoc analysis, all six primary intervals crossed zero. |
| Frozen game source | The evaluation was to use the designated frozen audit game source. | That source was missing at runtime, and the existing copy of the paired project was frozen in practice. The historical scenarios were restored afterward, and their fingerprints were checked. |
| Observation at the cutoff game tick | The legacy landmark interface was to be used to read the state at game tick 60. | The runtime environment provided observe, and the Python adapter combined it with the per-tick trajectory to reconstruct the observation at the cutoff. The endpoint was subsequently rechecked using the native observer. |
| Scenarios and receipts | The design identifier and scenario verification file were to be locked before unblinding. | The reconstructed scenario verification file did not match the originally registered hash. After evaluation, the scenario-verification fingerprint was updated, as were the design identifiers in 18,720 receipts. |
| Scope of rewriting | We compared the versions of each receipt for which an independent earlier copy was available. | Of the 18,720 relabeled receipts, 99 had independent early copies available for comparison, covering 3/9 combinations of scenario and input group. For the low-health start, the all-zero group had 81 per-episode receipts and 1 evaluation-freeze receipt. For the severe-thirst start, the intact group had 10 per-episode receipts. For the severe-hunger start, the shuffled group had 7 per-episode receipts. The comparable receipts differed only in the design identifier. All 98 earlier per-episode receipts came from pilot seeds. The 24 additional seeds had no comparable early per-episode receipts. The remaining 18,621 receipts lacked early copies. It remains to be verified that, before unblinding, a single plan was in place and the full set had been sealed. |

On each of the two days on chip 22021, one organoid was recorded, and each organoid had only one acquisition session. For the second specimen, the adverse-start trials replayed multiple H5 files from the same one acquisition session and involved 96 models, 32 seeds, and 64 maps. The comparison across five groups over the two days also used a common set of 8 seeds and 48 maps. The recordings, fixed readout panel validation, input structure, and trial scheduling for chip 22554 differed from those for chip 22021. On chip 22554, the intervals for all three exploration comparisons at six decisions crossed zero. The scope of these comparisons was limited to recordings from that chip and the controls tested. For the first specimen’s 16 training seeds and 48 held-out maps, we computed the difference in the number of newly revealed cells at each matrix entry by subtracting the all-zero group’s count from the intact or unstimulated group’s count for the same seed and map. For each comparison, we partitioned the total sum of squared deviations from the mean into seedrow, map-column, and within-cell residual components. At the end of six decisions, the proportions for the intact group were 2.44%, 29.10%, and 68.46%, and those for the unstimulated group were 1.26%, 38.58%, and 60.16%, in the order of seed rows, map columns, and within-cell residuals. For the same batch of trajectories through game tick 6, the corresponding proportions were 2.44%, 25.61%, and 71.96% for the intact group and 1.36%, 41.98%, and 56.66% for the unstimulated group. These proportions describe the matrices measured. The training seeds in the initial and expanded batches did not overlap. Seeds were therefore nested within batches, and the seed-row proportion included both within-batch seed differences and differences between the means of the two batches. There were no replicate games within any group at the same seed and map, so the within-cell residual could not further distinguish the seed-by-map interaction from other within-game variation. Variation across independent organoids, chips, and sessions cannot yet be estimated separately.

On chip 22021, there were only six rounds per day for each stimulation condition. On 11/19, sites H, W, and F on chip 22554 each had one screening stimulation recording for comparison of the preand poststimulation windows. Same-site zero-output controls were unavailable. After the fixed readout panel failed validation on 11/18, the exploratory acquisition was registered before it began as having received no approval for real-time training. Post-acquisition exploratory leave-one-block-out classification of the nine trials yielded 5/9 correct classifications both before and after stimulation (post-stimulation p*≈*0.306). The game replay for that day used a separate split into six training trials and three held-out trials, as described in Section S4.2. The total number of correct classifications after stimulation and the individual counts for sites H and F were all below the reference criteria in the qualification procedure. The available evidence supports engineering verification of commands and readback. The separability of neural responses by stimulation site and the actual waveform under electrode load remain undetermined.

The pre-registration deviation table lists the distribution of earlier copies by scenario and group. The comparable per-game receipts involved all 64 maps, but covered only a subset of scenario and input group combinations for the pilot seeds.

All 64 evaluation maps and 8 of the 32 training seeds had been used in the pilot. The post hoc sensitivity analysis retained only the remaining 24 seeds but still used all 64 pilot maps. It could assess only how sensitive the results were to excluding the pilot seeds. Independent validation on new maps remained unavailable. The table below presents the full sample and this subset side by side using the same direction of estimation. Each column’s intervals and counts of seeds with favorable differences were calculated using that column’s inference sample. For all six primary and four secondary comparisons, the intervals crossed zero in each of the two analyses, leaving the corresponding effects on exploration, survival, or waiting for supplies inconclusive.

### Full-sample analysis and sensitivity analysis using additional seeds for adverse starts

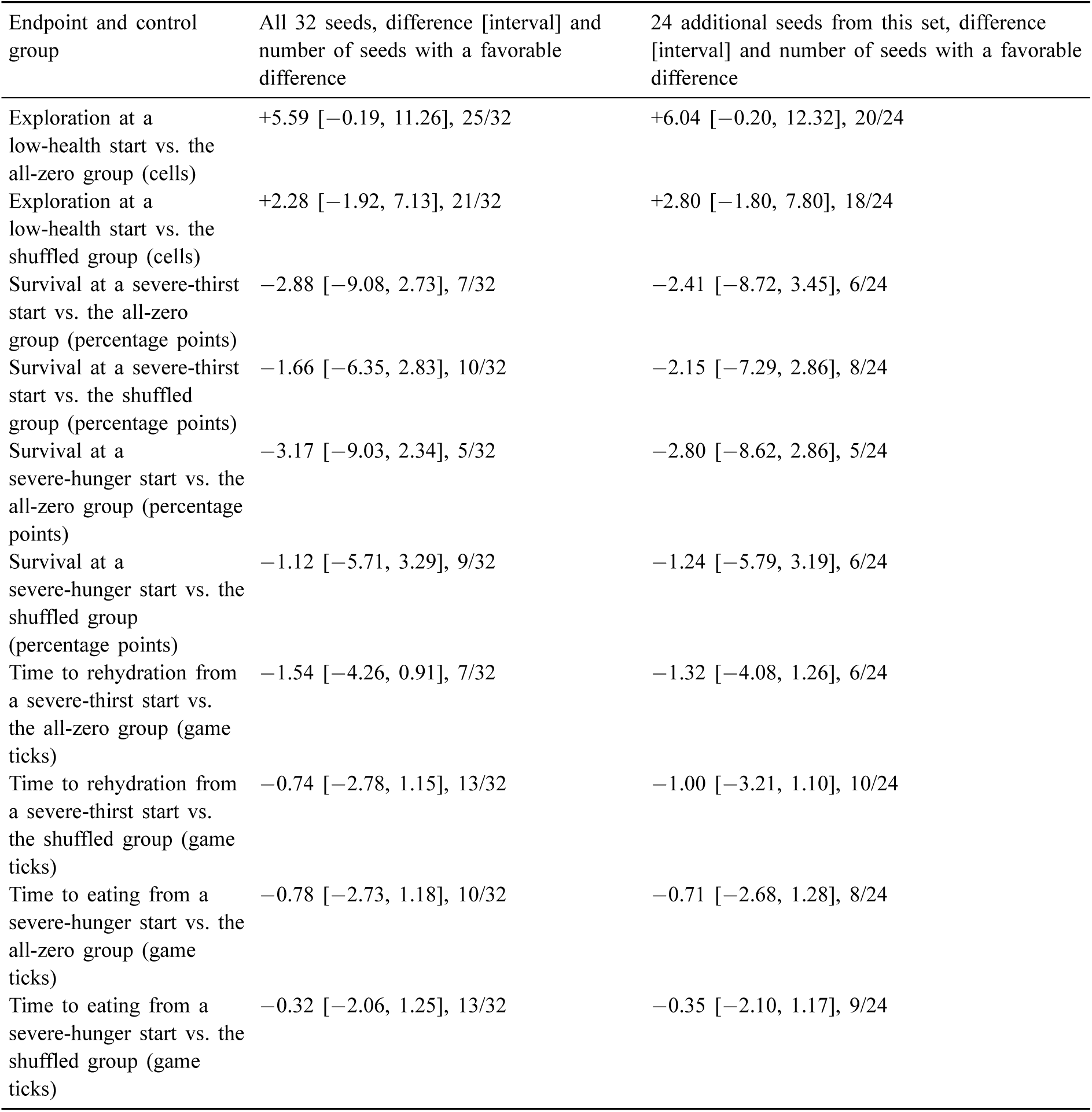

| Endpoint and control group | All 32 seeds, difference [interval] and number of seeds with a favorable difference | 24 additional seeds from this set, difference [interval] and number of seeds with a favorable difference |
|---|---|---|
| Exploration at a low-health start vs. the all-zero group (cells) | +5.59 [−0.19, 11.26], 25/32 | +6.04 [−0.20, 12.32], 20/24 |
| Exploration at a low-health start vs. the shuffled group (cells) | +2.28 [−1.92, 7.13], 21/32 | +2.80 [−1.80, 7.80], 18/24 |
| Survival at a severe-thirst start vs. the all-zero group (percentage points) | −2.88 [−9.08, 2.73], 7/32 | −2.41 [−8.72, 3.45], 6/24 |
| Survival at a severe-thirst start vs. the shuffled group (percentage points) | −1.66 [−6.35, 2.83], 10/32 | −2.15 [−7.29, 2.86], 8/24 |
| Survival at a severe-hunger start vs. the all-zero group (percentage points) | −3.17 [−9.03, 2.34], 5/32 | −2.80 [−8.62, 2.86], 5/24 |
| Survival at a severe-hunger start vs. the shuffled group (percentage points) | −1.12 [−5.71, 3.29], 9/32 | −1.24 [−5.79, 3.19], 6/24 |
| Time to rehydration from a severe-thirst start vs. the all-zero group (game ticks) | −1.54 [−4.26, 0.91], 7/32 | −1.32 [−4.08, 1.26], 6/24 |
| Time to rehydration from a severe-thirst start vs. the shuffled group (game ticks) | −0.74 [−2.78, 1.15], 13/32 | −1.00 [−3.21, 1.10], 10/24 |
| Time to eating from a severe-hunger start vs. the all-zero group (game ticks) | −0.78 [−2.73, 1.18], 10/32 | −0.71 [−2.68, 1.28], 8/24 |
| Time to eating from a severe-hunger start vs. the shuffled group (game ticks) | −0.32 [−2.06, 1.25], 13/32 | −0.35 [−2.10, 1.17], 9/24 |

For newly revealed cells and survival, the table reports the intact group minus the control group, with 99.167% intervals. For waiting time, it reports the control group minus the intact group, with 98.75% intervals. All four point estimates of waiting-time differences were negative. In both columns, runs in which death occurred before the first success or no success occurred by the cutoff were counted as 60 game ticks. In the separate crossed resampling for each column, the arms shared map indices. If the shuffled group was involved in the crossed resampling, the specified permutation was resampled along with the map. The additional-seed column presents a post hoc analysis that reused all 64 pilot maps.

## S9 Conclusion

In the offline replay of recordings from the first organoid recorded on chip 22021, under the same budget of six decisions, the intact group and the unstimulated group revealed an average of 1.99 and 2.16 more cells per run than the all-zero group, respectively. The mean differences were positive for 14/16 and 15/16 seeds, respectively. When runs were paired by training seed and map, the median difference was zero in each of the two comparisons. Cell counts tied in 615/768 pairs for the intact group comparison and in 602/768 pairs for the unstimulated group comparison. The number of newly revealed cells was identical across all three groups in 577/768 seed–map combinations. Of the 48 maps tested, 14 had identical cell counts across the three groups for all 16 seeds at the end of six decisions. This count was 20 when the same trajectories were truncated post hoc at game tick 6. At game tick 6, the mean differences for the intact group and the unstimulated group relative to the all-zero group were +1.21 and +1.53 cells per run, respectively. The archived per-comparison 95% crossed resampling intervals were both above zero. In a sensitivity analysis using the original *β* sweep’s nominal 98.333% level, the intervals for the two comparisons were all above zero when seeds were resampled on the fixed map set. The intervals for the two comparisons all crossed zero under crossed resampling of seeds and maps. All these intervals for game tick 6 used the complete data from all 48 maps as their starting point. Maps M29 and M45 were selected post hoc based on the observed net differences and were excluded together. All 16 training seeds were retained on the remaining 46 maps, giving 736 pairs per comparison. At game tick 6, the mean differences in newly revealed cells for the intact group and the unstimulated group relative to the all-zero group fell to +0.639 and +0.663 cells per run, respectively. Among the original 48 maps, at game tick 6, the map-level mean differences for the intact group and the unstimulated group relative to the all-zero group were positive on 17 and 16 maps, respectively, and negative on 8 maps for each comparison. The median of the map-level mean differences was zero in each of the two comparisons. The joint exclusion analysis described the differences on the retained maps. The difference in newly revealed cells at equal elapsed time on a different map set remained uncertain. Future work could independently validate the result on maps that were outside this analysis. Before inspecting the evaluation results, it could fix the count of newly revealed cells at game tick 6 as the exploration endpoint, along with the family of comparisons taking each group with measured inputs (intact and unstimulated) minus the all-zero group, the correction rules, and the criterion for stable improvement. Runs could be paired by training seed and map, with crossed resampling of seeds and maps. The overall paired mean differences, their intervals, and the distribution of map-level mean differences could then be reported. After pooling the two batches, the mean R1 differences at the end of six decisions for the intact group and the unstimulated group relative to the all-zero group were +0.004065 and +0.004339 points per run, respectively. The per-comparison 95% intervals from resampling seeds on the fixed map set were both above zero. When the same trajectories were truncated post hoc at game tick 6, the two mean R1 differences were +0.000443 and +0.000220 points per run, respectively. The per-comparison 95% intervals from crossed resampling of seeds and maps both crossed zero. The two score analyses differed in both the readout time and the resampling units. The timing decomposition was limited to the first day’s five-group subexperiment (8 seeds *×* 48 maps). All groups survived to the sixth decision. The timing term corresponding to cumulative action duration accounted for 84.3% and 95.9% of the two total R1 differences, respectively. The post hoc 97.5% intervals for the two residuals after subtracting the timing term all crossed zero. These differences were limited to the tested recordings and maps for the first organoid recorded on chip 22021. Each group used its own readout method during both training and inference. The design could neither separate the contribution of time-varying input in general from the contribution specific to its organoid origin nor distinguish action preferences learned during training from immediate responses at inference time.

The six-decision comparison for 22554 and the reanalysis of the same trajectories at game tick 6 used records from 11/18. The replay and control designs differed from those for 22021. After six decisions, the individual 95% intervals on the fixed map set for all three comparisons of newly revealed cells crossed zero. The intact group (ordered training pairings) and the shuffled group (shuffled training-trial pairings) scored 0.003388 and 0.003356 points per episode higher than the native R1-only group, respectively. Both corresponding intervals were above zero. These two score differences corresponded to the full set of integration conditions, which included adding four-dimensional MEA readouts and changing the value network input from 108 to 112 dimensions. The independent contributions of these changes could not be separated. Under shared ordered replay, the interval for the score difference between the intact group and the shuffled group, computed as intact minus shuffled, crossed zero. The same trajectories were retrospectively truncated at game tick 6. The individual 95% intervals from crossed resampling for all three differences in newly revealed cells and all three score differences crossed zero. A separate offline evaluation on 11/19 held out files by site. All four individual 95% intervals for differences in newly revealed cells at game ticks 48 and 192 crossed zero (Appendix B). For 22021, on each of the two days, the post-regular-stimulation group and the post-random-stimulation group were each compared with the unstimulated group. The archived adjusted intervals for differences in newly revealed cells and scores all crossed zero, and the effects of the tested stimulation conditions remained inconclusive. The second organoid recorded on chip 22021 was reassessed on all 64 reused pilot maps according to the available registration draft. None of the six primary comparisons met the criteria (0/6). In the low-health start, 2,048 of 2,048 episodes in each group survived to game tick 60, and the survival rate reached its ceiling. The primary endpoint for this start was the number of newly revealed cells at game tick 60 or at death if it occurred earlier. The intact group revealed 5.59 and 2.28 more cells per episode than the all-zero group and the shuffled group (whose two-second blocks were reordered within trials), respectively. The two adjusted 99.167% intervals crossed zero. At game tick 60, the differences in survival rates for the intact group minus the all-zero group and the intact group minus the shuffled group were *−*2.88 and *−*1.66 percentage points, respectively, in the severe-thirst start, and *−*3.17 and *−*1.12 percentage points, respectively, in the severe-hunger start. All four adjusted 99.167% intervals crossed zero. Across all episodes, the point estimates for the differences in restricted waiting time to drink and to eat (control group minus intact group) were all negative. All four adjusted 98.75% intervals crossed zero. Survival and restricted waiting time describe observable in-game behavior. Subjective fear was not measured. Future work could prespecify counts of newly revealed cells after six decisions and at game tick 6, along with survival at game tick 60, as endpoints using independent organoids, chips, and sessions, and maps unused in the pilot. The originally proposed 6 organoids, at least 3 chips, seed counts of 32, 64, and 96, and new-map counts of 64 and 128 were all candidate sample sizes. With 80% as the target power, future work could estimate the required sample size separately for each endpoint using its prespecified success criterion and biological-level variability. It could also estimate the sample size needed to meet the criteria for multiple endpoints jointly.

## Appendix A Per-seed results

Table 5 lists the total number of newly revealed cells across 48 games for each training seed of the first organoid recorded on chip 22021. Summing the rows for an arm and dividing by 768 games yields the pergame mean. Compared with the all-zero group, the intact group had higher counts for 14 seeds and lower counts for 2 seeds. The unstimulated group had higher counts for 15 seeds and the same count as the all-zero group for 1 seed. All training seeds shared the same maps.

**Table 5.** Total newly revealed cells across 48 games for each training seed. Each of the two difference columns uses the all-zero group in the same row as the control. All three groups share the same maps. Different games can reveal the same location separately, so the cumulative counts include overlapping cells.

| Training seed | All-zero group | Intact group | Unstimulated group | Intact group − all-zero group | Unstimulated group − all-zero group |
|---|---|---|---|---|---|
| 102083199 | 1,869 | 1,983 | 1,927 | +114 | +58 |
| 102122616 | 1,926 | 2,079 | 2,125 | +153 | +199 |
| 102385481 | 2,031 | 2,212 | 2,233 | +181 | +202 |
| 102411608 | 1,937 | 2,106 | 2,069 | +169 | +132 |
| 102452542 | 1,968 | 1,958 | 1,993 | −10 | +25 |
| 102473463 | 1,961 | 1,904 | 2,071 | −57 | +110 |
| 102476656 | 1,885 | 1,972 | 2,053 | +87 | +168 |
| 102481912 | 1,969 | 2,030 | 2,011 | +61 | +42 |
| 102600001 | 1,909 | 1,928 | 1,985 | +19 | +76 |
| 102600002 | 1,938 | 2,134 | 2,019 | +196 | +81 |
| 102600003 | 1,991 | 1,997 | 1,991 | +6 | 0 |
| 102600004 | 2,034 | 2,131 | 2,162 | +97 | +128 |
| 102600005 | 1,901 | 1,977 | 2,027 | +76 | +126 |
| 102600006 | 1,952 | 2,155 | 2,017 | +203 | +65 |
| 102600007 | 1,875 | 1,948 | 1,995 | +73 | +120 |
| 102600008 | 1,871 | 2,029 | 1,996 | +158 | +125 |

In a post hoc analysis, we excluded each map in turn along with all seed records for that map. For each exclusion, we calculated the mean difference from 752 pairs of games on the remaining 47 maps. At the end of six decisions, the mean difference ranged from +1.479 to +2.207 cells per game for the intact group minus the all-zero group and from +1.512 to +2.439 cells per game for the unstimulated group minus the all-zero group. At game tick 6 in the same trajectories, the corresponding ranges were +0.924 to +1.399 and +1.074 to +1.779 cells per game, respectively. The minima for the first three ranges occurred when M29 was excluded, and the minimum for the last range occurred when M45 was excluded. These ranges describe how the descriptive point estimates varied with the composition of the tested map set.

Supplementary Table. Per-map mean differences in newly revealed cells for the first specimen. Each table cell gives the mean paired difference in newly revealed cells across 16 training seeds on the same map (cells per game).

| Anonymous map | After six decisions · intact minus all-zero | After six decisions · unstimulated minus all-zero | At game tick 6 · intact minus all-zero | At game tick 6 · unstimulated minus all-zero |
|---|---|---|---|---|
| M01 | 0.9375 | 0 | 4.375 | 3.5 |
| M02 | 9.9375 | 6.625 | 4.5 | 3 |
| M03 | 5.75 | 4.3125 | 5.75 | 4.3125 |
| M04 | 0 | 0 | 0 | 0 |
| M05 | 0 | 0 | 0 | 0 |
| M06 | 0.3125 | 0 | 0.625 | 0 |
| M07 | 0 | 0 | 0 | 0 |
| M08 | 0 | 0 | 0 | 0 |
| M09 | −8.375 | −11.0625 | −7.4375 | −10.125 |
| M10 | 4.6875 | 4.6875 | 0 | 0 |
| M11 | 0 | 0 | 0 | 0 |
| M12 | 0 | 0.3125 | 0 | 0 |
| M13 | −0.9375 | 1.875 | −0.9375 | 1.875 |
| M14 | −0.375 | −0.75 | −0.375 | −0.75 |
| M15 | 3.75 | 2.1875 | 0 | 0 |
| M16 | 1.5625 | 1 | 0 | 0.0625 |
| M17 | 0 | 0 | 0 | 0 |
| M18 | −0.9375 | 3.625 | 2.375 | 8.5 |
| M19 | 1.375 | −0.75 | 0 | 0 |
| M20 | 6.875 | 7.125 | 2.75 | 4.6875 |
| M21 | 0 | 0 | 0 | 0 |
| M22 | 3.5 | 3.625 | 3.5 | 4.5 |
| M23 | 1.75 | 0.875 | 1.75 | 0.875 |
| M24 | 0 | 0 | 0 | 0 |
| M25 | 16 | 13.625 | 10.5 | 9 |
| M26 | −3.75 | −3.75 | −2.5 | −2.5 |
| M27 | −0.625 | −0.75 | −0.625 | −0.75 |
| M28 | −1.625 | −1.625 | −1.75 | −1.75 |
| M29 | 25.875 | 32.5 | 14.875 | 20 |
| M30 | 0 | 0 | 0 | 0 |
| M31 | −1.875 | −0.9375 | −1.875 | −0.9375 |
| M32 | 12.5 | 12.25 | 3.125 | 3.6875 |
| M33 | 2.625 | 3.875 | 2 | 3.25 |
| M34 | 0 | 0 | 0 | 0 |
| M35 | 1.4375 | 4.8125 | 1.4375 | 3.1875 |
| M36 | 0 | 0.25 | 0 | 0.25 |
| M37 | −1.875 | −3.75 | 0 | 0 |
| M38 | 1.875 | 2.8125 | −2 | −3 |
| M39 | 2.5 | 1.5 | 0 | 0 |
| M40 | 0 | −0.75 | 0.375 | −0.375 |
| M41 | 0.125 | 0 | 2.875 | 0 |
| M42 | 0 | 0 | 0 | 0 |
| M43 | 0 | 0 | 0 | 0 |
| M44 | 0 | 0 | 0 | 0 |
| M45 | 13.6875 | 22.0625 | 14.0625 | 23 |
| M46 | −2.25 | −2.25 | 0 | 0 |
| M47 | 0.9375 | 0 | 0.9375 | 0 |
| M48 | 0 | 0 | 0 | 0 |

## Appendix B Reproduction and quality control

According to the consolidation register for the retained evaluation logs, the six-decision trial on the first organoid recorded on chip 22021 included 384 games per arm in each batch. Across two batches, there were 768 games per arm and 2,304 games across three arms. Games with the same map and action sequence were consolidated into 146 distinct combinations. A complete anonymized matrix of paired per-game counts of newly revealed cells and an accompanying program for recomputation are also available. The historical launcher for the first organoid’s trial pointed to Godot 4.7.2 for Linux. The independent replay used Godot 4.7.2.stable.official.ed1daf0bf for Windows. The scene came from later copies of the project for the same scene that matched file for file across the runtime archives from two nodes. The replay submitted the 876 saved actions from the 146 combinations to the engine and mapped the results back to the corresponding entries for all 13,824 retained decisions from two batches and three groups. Post-action states and diagnostic R2 labels matched at every step, and the maximum absolute difference in stepwise R1 was zero. Across all 2,304 games, the number of newly revealed cells, the final game tick, health, hydration, satiety, and termination flags matched game by game. The maximum absolute difference in end-of-game R1 was approximately 2.78*×*10*^−^*^17^ points. Pre-action game observations were encoded before the MEA readings were added. We sorted the output of encode(obs, ’reserves’), serialized it as compact JSON, encoded it in UTF8, and computed its SHA-256 digest for comparison with the rows[].context field of the retained decisions.

When we recomputed the digests using native Windows floating-point values, 2,358 retained digests matched and 11,466 differed. For each of the three resources, wood, stone, and food, we added one to the count, took the natural logarithm, and divided by five. Using integer resource counts as keys, we looked up the archived floating-point values for these three features from 1,152 decisions in the 96 training logs of the first batch. For a retained first step with a food count of 2, we replaced only the food feature at features[5] in the base game feature vector, substituting the archived training value 0.21972245773362192 for the Windows value 0.21972245773362195. The digest for that step then matched. The counts.holdout_training_precision_ equal field in the diagnostic summary recorded that all 13,824 retained digests matched after recomputation at the archived precision. The counts.training_files, counts.training_decisions, and counts.training_native_ mismatches fields in that summary recorded, respectively, the number of training logs in the first batch, the number of decisions, and the number of decisions with native numerical discrepancies. For 856 of the 1,152 training decisions, features recomputed using native Windows values differed from the archived training values. The effect on model outputs remains to be checked. The replay verified game outcomes for the supplied actions and the encoding of game observations before MEA readings were added. Evidence is still missing both for independent reproduction of batch-by-batch training updates and for independent regeneration of actions after reloading the frozen policy.

In the adverse-start trial using the second organoid on chip 22021, we re-extracted signals for 324 channels from the 19 included H5 files. We verified 24 calibration windows, 48 training windows, 4 subsequent valueestimation windows, and 60 held-out evaluation windows. We also checked 32 training permutations and 64 evaluation permutations. For each of the two organoids on chip 22021, we fitted the calibration parameters using 24 consecutive one-second windows from its own warm-up recording for the work reported here. After conversion from device frames, the first specimen’s calibration windows covered [12, 36) seconds after warm-up. They partially overlapped the training windows of the intact group in the first-day five-group comparison described in Section S4.3 and were separate from that group’s held-out windows at [58, 64) seconds after warm-up. The two interfaced groups in the 22554 six-decision trial used calibration parameters from 50 windows in the archived baseline. We checked the replay files and training records for those groups. On 11/19, the 22554 trial used the alternative windows described in Section S4.1 to conduct offline comparisons over 48 and 192 game ticks. The plan and windows fields in the interfaced groups’ training records contain the file fingerprints for the alternative screening protocol and the window list, respectively. The hold field lists the held-out sites. Each of the three folds, H, W, and F, held out one site file. We removed commonmode transients separately for each site. In each fold, we fitted the four-dimensional standardization parameters using only one prestimulus window from each of the remaining two sites. For every site, including the held-out site, we selected readout channels using the pre- and post-stimulus windows from that site’s own file. Held-out evaluation was conditional on obtaining and preprocessing that file. The archived screening program used voltage and event times. The effect of this preprocessing on game outcomes remains unquantified. The following day’s leave-one-file-out entry in the per-model budget inventory showed that 12 interfaced models were newly trained that day, each completing 96 training decisions. The held-out evaluation of another 6 native R1-only models reused existing checkpoints with matching fingerprints and used only native game observations. The earlier training summaries each recorded 96 decisions and 8 updates. The comparability of the two training setups across the full training process remains to be verified, as do the elapsed time on the original hardware and the GPU-hours. During training, the two interfaced groups used an exogenous auxiliary reward assigned by replay-window phase. The auxiliary term for post-stimulus windows used the negative sign associated with the stimulation record. In the training configuration, the R1 weight was 1, the R2 weight was 0.1, and the amplitude fraction was 0.5, corresponding to mix.r1_weight, mix.r2_weight, and mix.amplitude_fraction, respectively. All 1,152 checked training decisions across the 12 interfaced models recorded a nonzero auxiliary term. The max_weighted_r2_fraction field in the archived results indicated that, among decisions with nonzero game reward, the maximum absolute ratio of the weighted auxiliary term to the game reward was 5%. The intact-minus-shuffled contrast compared the two interface arrangements used during training. The contrast of the intact group minus the native R1-only group with reused checkpoints also involved differences in replay inputs, training arrangements, and auxiliary rewards. Their individual contributions still cannot be separated. The source-material directory inventory contained 183 entries, comprising 182 H5 files and the index file itself. The 192 rows in the old index corresponded to 182 unique filenames and 182 unique SHA-256 hashes. Of those filenames, 19 carried a confirmatory label. The actual scope of re-extraction for the adverse-start trial was based on the verification of trial sources described at the start of this paragraph. The file inventory consolidated duplicate sources by full hash and retained the old index’s flags for anomalies and confirmatory inclusion. A complete record of all on-site acquisition attempts and the rationale for each manual inclusion or exclusion decision is still missing. The step-by-step archive verification records showed that the training permutation was identical across training seeds in the shuffled group. In held-out evaluation, the permutation assigned to each map was reused across training seeds and start scenarios. At each step, the archived window ID, event fingerprint, and normalized readings matched the window assigned to that map at the corresponding game tick. The game ticks actually visited in an episode could vary with the selected actions. In the accompanying Input Files and Game Decisions Index, first locate the corresponding entry in the episode-level index confirmed_eval_traces.csv by training seed, start scenario, group, and map. In that table, archive_member and member_bytes_sha256 identify the archived trace member and its file fingerprint, respectively. The actual_window_ids field links, in decision order, to diagnostic_window_ id in the held-out portion of the per-window index confirmed_source_windows.csv. Use the window ID and device-frame range in the per-window index to look up the acquisition session and event fingerprint. Use raw_sha256, downloaded_cfg_sha256, roi_sha256, and destination_member_bytes_sha256 to verify the fingerprints of the raw H5 file, downloaded routing configuration, readout channel set, and archive member containing the four-dimensional features, respectively. The shuffled group’s permutations can be checked by map and game tick in confirmed_orders.csv. Training traces can be looked up by group, seed, batch, and training map in confirmed_training_traces.csv. The mapping from readout channels to physical electrodes is in roi_ electrodes.csv. For example, in the shuffled group under the severe-hunger start, the episode for training seed 351150800 on map 352002256 read window 1308 at its first step. Archive verification covered index-key joins for all held-out decisions. Independent row-by-row verification remains incomplete. The records of endof-batch value estimates include the designated windows and scalar outputs. Separate snapshots of the inputs actually read during the forward pass are still missing.

We read all 96 final optimizers.pt files from the adverse-start trial. For each model, we verified the step counts in the two sets of AdamW state. The policy took 8 steps and the value network took 64 steps, for totals of 768 and 6,144 steps, respectively. Fingerprint checks passed for all 576 final components. We reloaded each final model and reran it on the first map of each of the three scenarios. Across the three scenarios, there were 288 games in total, and 9,351/9,351 actions matched. The PEFT field in all 96 adapter configurations was 0.21.2. All 9 logs in which the engine version string was found contained a traceback or ERROR. We verified the adverse-start training updates against the batch-level logs and final optimizer states, the frozen policy reloads by comparing actions step by step on the first map, and the replay of saved actions against the complete set of game results. The fit.log file in the adverse-start training log bundle recorded successful jobs and their assigned GPU indices by training seed and group. The logs for successful jobs also recorded the PEFT and FLA call paths. The initial policy weight digest in the training registry matched the delivered warm-start policy, and the base configuration digest recorded in the final policy matched the delivered configuration. The verifier script digest and the action mismatch count for the frozen policy reload are recorded in policy-reload.json under verifier_sha256 and action_mismatches, respectively. The reload_trajectories.py script identified by that digest called torch.manual_seed to fix the random seed for the reload. It checked the initial state against the training registry, loaded the checkpoint and checked the frozen state, then compared actions step by step under torch.inference_mode(). The exit code for the replay of saved actions is recorded in native-replay.rc. The replay script and source receipt digests are recorded in native-replay.json under verifier_sha256 and source_receipt_sha256, respectively. The digests of the engine binary, transport code, and reward code used for that replay are recorded in native-sources.json. The delivered base weight shards can be verified against the file manifest. The base entries in the training update and frozen reload receipts record the frozen parameter tensor digests. The FLA and causal-conv1d file digests listed under kernel_identity in the final policy matched the manifest for the historical run bundle. The value network’s single-precision layout is recorded in the policy metadata. The engine digest for the historical action replay also matched the Linux engine file in that manifest. Complete per-process load manifests are missing for successful training runs and frozen reloads. The actual base files, interpreter, engine and full dependency set, GPU model, overall numerical precision, and determinism settings still cannot be verified individually after the fact. The Windows engine version mentioned earlier in the appendix applied only to the replay of saved actions from the first specimen’s six-decision trial. The original training wall-clock time and GPU-hours remain to be verified.

The power recalculation used a CPU, C++17, MSVC 19.44.35229.0, AVX2, 9 concurrent worker processes, and master random seed 2026112202. The coordinator recorded 3.453 seconds of wall-clock time for this invocation. Compilation and independent verification of the indices were accounted for separately. The recalculation did not train any models. An independent program checked 8 outer samples, and 80 threshold outcomes matched. To reproduce the analysis, we first formed pairs by pilot seed and map. We then used common resampling indices to compute the interval for each contrast and count the seeds with a difference in the positive direction. Finally, we counted the outer samples in which both contrasts succeeded for the same endpoint.

The Appendix B table “Next-day offline paired results for 22554 with files held out by site” reports results paired by training seed, held-out site, and shared map. There were 72 pairs per comparison and 72 games per arm. The two cutoff game ticks were evaluated using the same set of 216 games. According to the hold field in the training registry for the MEA-input group, the two groups shared the same mapping from seeds to held-out sites. Each seed corresponded to only one held-out site, and H, W, and F each had 2 seeds. Each fold yielded 24 pairs per comparison across the 12 shared maps. The native R1-only group reused existing checkpoints and received no MEA replay readouts at evaluation. See the earlier description of next-day training and preprocessing in this appendix. According to the archived analysis summary, each crossed resampling draw included 6 training seeds and 12 shared maps, with 20,000 draws in total. The two arms in each comparison shared the same indices. Square brackets indicate two-sided 95% intervals for each comparison. In the public reconstruction, seeds were sampled with replacement across the pooled training seeds. Each sampled seed carried its held-out site file, so the number sampled from each fold could vary. Map draws were shared across folds and groups, and the same draw indices were used for both time budgets. The recomputed intervals matched all 72 archived interval endpoints. The historical random-number generator state was not saved separately. The training files overlapped across folds. The intervals were conditional on the site files used in this analysis and their offline preprocessing. Variability due to new independent site files has yet to be estimated. The analysis was exploratory and used no adjustment for the family of endpoints. All four intervals for paired differences in the count of newly revealed cells crossed zero. For the intact group minus the native R1-only group, the intervals for the number of drinking events and the number of drinking events followed by a net gain in hydration were below zero at each of the two budgets. The interval for the difference in hydration reserves was below zero at game tick 48 and crossed zero at game tick 192. At game ticks 48 and 192, each of the three groups had 0/72 games with a death. At each cutoff game tick, each of the two paired differences in the proportion of games with a death was zero. Because there were no deaths in this batch, the empirical resampling intervals for these paired differences all collapsed to [0, 0]. Differences in the risk of death and equivalence on untested maps or new records remain undetermined.

Archive checks confirmed that the adverse-start trial for the second specimen of 22021 replayed 669,715 saved actions across 18,432 episodes using the replacement frozen game source. For every episode, the state-transition summary and the endpoints for exploration, survival, and restricted waiting time for supplies matched the archive. The final frozen policy was reloaded for the 288 episodes spot-checked above. Section S4.1 lists the pass counts and the real-time threshold for the three warm-up windows of the first specimen of 22021. For each site of 22554 on 11/19, it also lists the number of channels that passed all six windows, the threshold, and the window results before and after replacement. The warm-up pass rates for the first specimen of 22021 were checked during that acquisition session. All three windows fell below the real-time training threshold. The six-window results for 22554 came from offline screening after the 11/19 acquisition. At every site, the number of channels that passed all six windows fell below the screening threshold. The replacement windows for 22554 were used in the 11/19 offline comparisons at game ticks 48 and 192. Both the six-decision results and the recheck at game tick 6 for the same batch used the 11/18 replay.

### Next-day offline paired results for 22554 with files held out by site

| Budget and endpoint | Group means (in order: intact group, native R1-only group, shuffled group) | Intact group minus native R1-only group [interval] | Intact group minus shuffled group [interval] |
|---|---|---|---|
| 48 game ticks · newly revealed cells (cells per episode) | 108.46/108.14/108.40 | +0.32 [−19.97, 19.07] | +0.06 [−11.24, 12.31] |
| Game tick 48 · Drinking events (per episode) | 1.10/2.93/1.26 | −1.83 [−3.42, −0.49] | −0.17 [−0.99, 0.56] |
| Game tick 48 · Eating events (per episode) | 1.10/1.01/1.11 | +0.08 [−0.19, 0.36] | −0.01 [−0.26, 0.21] |
| Game tick 48 · Hydration (reserve points) | 83.32/89.59/85.06 | −6.28 [−12.07, −1.39] | −1.74 [−6.39, 2.80] |
| Game tick 48 · Satiety (reserve points) | 86.23/85.20/86.66 | +1.03 [−1.80, 4.45] | −0.43 [−2.70, 1.66] |
| Game tick 48 · Health (points) | 100.00/100.00/100.00 | 0 [0, 0] | 0 [0, 0] |
| Game tick 48 · Net increases in hydration after drinking (events per episode) | 0.75/1.65/0.85 | −0.90 [−1.67, −0.22] | −0.10 [−0.625, 0.31] |
| Game tick 48 · Net increases in satiety after eating (events per episode) | 1.07/0.97/1.07 | +0.10 [−0.14, 0.36] | 0.00 [−0.22, 0.21] |
| Game tick 48 · Cumulative mortality proportion | 0/72/0/72/0/72 | 0 [0, 0] | 0 [0, 0] |
| Game tick 192 · Newly revealed cells (cells per episode) | 243.03/277.28/238.46 | −34.25 [−76.25, 4.60] | +4.57 [−28.25, 38.81] |
| Game tick 192 · Drinking events (per episode) | 3.38/12.01/3.58 | −8.64 [−13.36, −4.50] | −0.21 [−1.51, 1.04] |
| Game tick 192 · Eating events (per episode) | 3.93/3.90/3.89 | +0.03 [−0.86, 0.88] | +0.04 [−0.53, 0.57] |
| Game tick 192 · Hydration (reserve points) | 87.51/93.07/89.51 | −5.56 [−11.26, 0.60] | −2.00 [−8.96, 5.19] |
| Game tick 192 · Satiety (reserve points) | 89.99/90.35/89.52 | −0.36 [−9.22, 7.22] | +0.47 [−3.80, 4.78] |
| Game tick 192 · Health (points) | 97.59/98.22/95.74 | −0.63 [−4.05, 2.57] | +1.85 [−0.59, 4.74] |
| Game tick 192 · Net increases in hydration after drinking (events per episode) | 2.51/6.60/2.67 | −4.08 [−6.47, −1.83] | −0.15 [−1.17, 0.85] |
| Game tick 192 · Net increases in satiety after eating (events per episode) | 3.78/3.64/3.71 | +0.14 [−0.65, 0.93] | +0.07 [−0.47, 0.57] |
| Game tick 192 · Cumulative mortality proportion | 0/72/0/72/0/72 | 0 [0, 0] | 0 [0, 0] |

Of the 18,432 episodes in the full replay, 4,167 had actions that crossed game tick 60. When R1 was recomputed at the game tick 60 boundary, the absolute difference from the archived total score exceeded 10*^−^*^10^ points in 64 episodes, with a maximum of 0.019450207 points. The primary analyses of adverse starts used exploration and survival at the cutoff. The secondary analyses used waiting time capped at game tick 60. The full-replay differences for these endpoints were zero. The six-decision scores and the first-day timing breakdown each used the archived R1 from the corresponding short-run trial. All 64 episodes had actions that crossed game tick 60. When the archived total score was subtracted from R1 recomputed at the game tick 60 cutoff, the difference was positive in 41 episodes and negative in 23. The absolute difference exceeded the threshold above in 3 low-health start episodes, 27 severe-thirst start episodes, and 34 severe-hunger start episodes. For each scenario and group, the signed difference was averaged across all 2,048 episodes. The largest absolute value of those means was approximately 0.00003454 points per episode. After switching to cutoff scoring, the largest absolute change in the paired mean difference between the intact group and either of the two controls across scenarios was approximately 0.00002775 points per episode. In the original evaluation, when an action crossed game tick 60, the Python adapter mapped the legacy observation request to observe and filled in some boundary fields from the last per-tick record at or before the cutoff. The remaining fields came from the observation at the end of the action. The retrospective replay read the complete native observation saved by the game at game tick 60. Both scoring methods computed the discounted timing term using the retained game ticks and computed the potential term using their respective boundary observations. In the per-episode score verification table native-replay.csv, r1_native stores the native score recomputed at the cutoff, and r1_recorded stores the archived total score. The signed difference was computed by subtracting the latter from the former. The group table r1_by_scene_arm.csv lists summaries for nine scenario-bygroup combinations. The table r1_comparison_sensitivity.csv lists six changes in paired mean differences by scenario and control, with the corresponding number of pairs for each comparison. The archived verification summary reported that the per-episode recomputations agreed with both detailed tables entry by entry. The public episode-level cutoff scores reproduce all nine group summaries and six paired contrast changes by decimal arithmetic. The absolute value of the mean signed difference was largest for the intact group in the severe-thirst start scenario. The absolute change in the paired mean difference was largest for the intact group minus the all-zero group in that scenario. The original run did not save the two types of complete boundary observation as pairs for each action, so the specific fields underlying each above-threshold difference remain to be verified. Section S8 checks registration against execution and details the replacement of the frozen source, the adapter, and the changes in scenario hashes.

Table 6 lists the training hyperparameters. The policy and value networks were updated separately, with learning rates of 0.000002 and 0.0001, respectively. The PPO clipping parameter was 0.1, the target KL was 0.015, and the per-time-step discount was 0.99985, with *α*=1 and *β*=0. Each model was trained for 4 batches of 12 decisions each, for a total of 48 training decisions, as described in Algorithm 1 and Table 6. The frozen evaluation for adverse starts ended at game tick 60 or at death if it occurred earlier, as described in Algorithm 2.

**Table 6.** Training settings for the adverse-start trial of the second specimen of 22021. Training comprises 48 decisions per model, with 32 models for each of the three input types.

| Parameter | Value | Purpose |
|---|---|---|
| Number of training decisions and optimization steps | 4 batches × 12, with 8 Actor updates and 64 Critic updates | Training used the same number of interaction records in each batch. |
| Learning rate / training epochs per batch | Actor 0.000002 / 2; Critic 0.0001 / 16 | Controlled the step size and number of repetitions for updates to the two sets of parameters. |
| PPO clipping / reference constraint | 0.1 / 0.02, target KL 0.015 | In the PPO surrogate loss, the archived code clipped the ratio of the joint probability of the selected verb and target under the current policy to that under the data-collection policy. The reference term used the KL divergence from the current policy to the archived reference distribution. The conditional target term was weighted by the ratio of the selected verb's probability under the current policy to that under the data-collection policy. The entire reference KL term in the policy loss was then multiplied by 0.02. When the approximate KL divergence from the data-collection policy to the current policy exceeded 0.015, early stopping occurred before the policy update for that step was applied. |
| Time-step discount / advantage decay | 0.99985 / 0.999 | Determined the weight of future scores in the current estimate. |
| Action verb exploration probability / entropy coefficient / reward weight | *ϵ*=0.1 / 0.002 / *α*=1, *β*=0 | The exploration probability and entropy coefficient controlled action exploration and policy entropy regularization, respectively. Only the game score was used as the training reward. |

## Data and code availability

The game and training code, paired outcomes, and archive manifests are available in the project repository at https://github.com/brainintelli/BioSignal-Guided-LLM-Reinforcement-Learning. The data access guide gives instructions for obtaining and verifying the raw H5 recordings.

## Use of AI tools

OpenAI Codex assisted with English translation and back-translation, editorial revision, preparation of analysis and figure-generation scripts, and checks of numerical results against archived outputs. The Qwen policy evaluated in the survival game is described in Section S3.3.

## Authors and contributions

Haoman Chen, Qingzu He, Fanxuan Chen, Xinyu Chen, Yang Liu, and Pengcheng Yan contributed equally, regardless of listing order.

Jianwei Shuai is the corresponding author.

